# Diverse dynamics of glutamatergic input from sensory neurons underlie heterogeneous responses of olfactory bulb outputs *in vivo*

**DOI:** 10.1101/692574

**Authors:** Andrew K. Moran, Thomas P. Eiting, Matt Wachowiak

**Affiliations:** Interdepartmental Program in Neuroscience, University of Utah School of Medicine, Salt Lake City, UT; Department of Neurobiology and Anatomy, University of Utah School of Medicine, Salt Lake City, UT

**Keywords:** imaging, glutamate, calcium, temporal coding, active sensing

## Abstract

Mitral/tufted (MT) cells of the olfactory bulb (OB) show diverse temporal responses to odorant stimulation that are thought to encode odor information. To understand the role of sensory input dynamics versus OB circuit mechanisms in generating this diversity, we imaged glutamate signaling onto MT cell dendrites in anesthetized and awake mice. We found surprising diversity in the dynamics of these signals, including excitatory, suppressive, and biphasic responses as well as nonlinear changes in glutamate signaling across inhalations. Simultaneous imaging of glutamate and calcium signals from MT cell dendrites revealed highly correlated responses for both signals. Glutamate responses were only weakly impacted by blockade of postsynaptic activity, implicating sensory neurons as a primary source of glutamate signaling onto MT cells. Thus, the dynamics of sensory input alone, rather than emergent features of OB circuits, may account for much of the diversity in MT cell responses that underlies OB odor representations.

## INTRODUCTION

The neural activity underlying odor perception is inherently dynamic. In vertebrates, olfactory sensory input arrives at the brain as bursts of activity driven by each inhalation, and patterns of inhalation are actively controlled by an animal as it samples its environment. At the level of the olfactory bulb (OB), olfactory sensory neurons (OSNs) expressing a single type of odorant receptor converge onto glomeruli^1^ where they make glutamatergic synapses onto targets including juxtaglomerular interneurons and the principal OB output neurons, mitral and tufted (MT) cells^2^. Odorant responses among MT cells can be temporally complex, displaying sequences of excitation and suppression that occur relative to a single inhalation and across multiple inhalations^3-6^. Olfactory information is likely represented both by the identity of activated MT cells as well as by their temporal dynamics^4,7-9^.

Much of the complexity in MT cell response patterns has been attributed to processing by inhibitory OB circuits^10-13^. However, excitatory synaptic inputs to MT cells, which occur solely on the MT cell apical tuft in each glomerulus, can also contribute to generating temporally complex and diverse patterns of MT cell spiking. OSNs themselves respond to odorant inhalation with different latencies and show nonlinear changes in their activation patterns as a function of odorant concentration, sampling time and sampling frequency^14,15^. Additional diversity in excitatory synaptic input to MT cells may arise from multisynaptic pathways – for example, via external tufted (ET) cells^12,16-18^. However, the contribution of glutamatergic input dynamics to determining MT cell odorant response patterns in vivo remains poorly understood.

Here, we imaged odorant-evoked and inhalation-linked glutamate signals in glomeruli of the mouse OB using the genetically-encoded glutamate sensor iGluSnFR and its second-generation variants^19,20^, expressed selectively in MT cells. Glomerular glutamate transients reported inhalation-linked dynamics with high fidelity, allowing temporally precise monitoring of glutamatergic signaling across the dorsal OB in both the anesthetized and awake mouse. We observed a striking degree of diversity in the temporal dynamics of glutamatergic input onto MT cell dendrites in a glomerulus, over time scales involving a single inhalation and across multiple inhalations of odorant. We found a high correspondence between the dynamics of glutamatergic input and that of MT cell output across glomeruli. Glutamate responses were only weakly impacted by blockade of postsynaptic activity, pointing to OSNs as the primary source of odorant-evoked MT cell excitation and a subtle role for multisynaptic excitation. These results reveal remarkable diversity in the dynamics of glutamatergic drive onto MT cells and suggest that the complex spatiotemporal activity patterns thought to play a critical role in coding odor information may arise largely from diverse patterns of feedforward excitation driven directly by sensory afferents.

## RESULTS

### Diverse inhalation-linked glutamate transients imaged from MT cell dendrites

The timing of odorant- and inhalation-evoked MT cell spiking varies across MT cells and across odorants, and has long been hypothesized to play a role in coding odor information^4,7,8^. To assess the degree to which such variability is present at the level of glutamatergic input to MT cells, we targeted iGluSnFR and its second-generation variants to MT cells using Cre-dependent viral vectors and MT cell-specific Cre driver lines^21,22^ (Fig. 1a). We have previously shown, using the first-generation iGluSnFR, that each inhalation of odorant drives a transient of glutamate release onto juxtaglomerular interneuron processes in the glomerular neuropil^23^. We observed similar glutamate signals with iGluSnFRs expressed in MT cells, with different glomeruli showing glutamate transients with varying onset latencies and durations, and with distinct changes in the glutamate signal evoked by successive inhalations across a longer odorant presentation (Fig. 1b). In the superficial OB layers, glutamate signals appeared restricted to MT cell apical tufts within glomeruli, with negligible signals outside of the glomerulus or on MT cell primary dendrites or somata (Fig. 1c, d). Odorant-evoked glutamate signals were also detected along MT cell lateral dendrites in the external plexiform layer (Supplementary Fig. 1a, b).

**Fig. 1.**
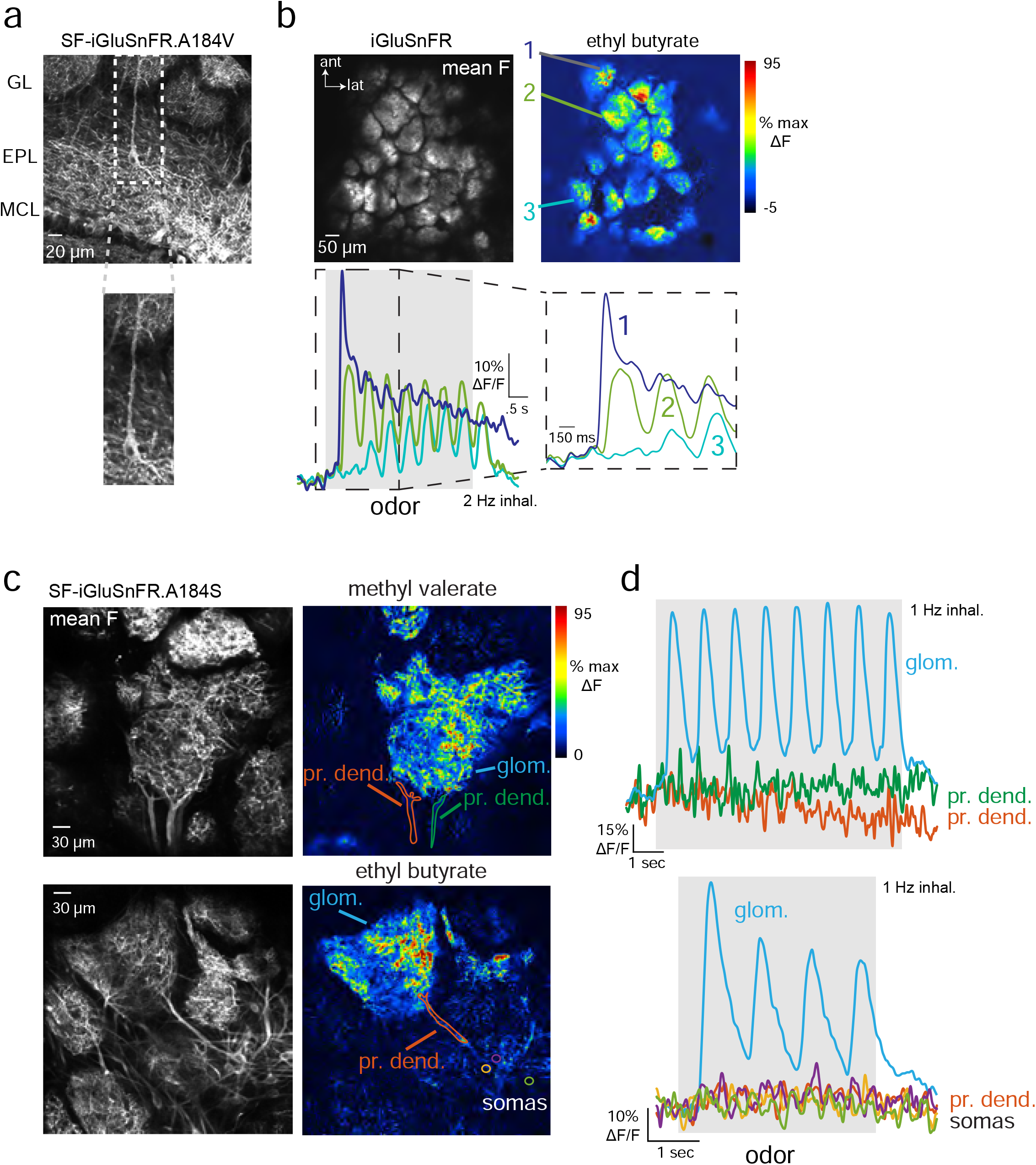
Characterization of iGluSnFRs as reporters of glutamatergic signaling onto MT cells. **a.** Expression of SF-iGluSnFR in MT cells of the OB. Image shows confocal stack with expression in MT cells after AAV2.1.hSynap.Flex.SF-iGluSnFR.A184V in the OB of a Tbet-Cre mouse. White arrows indicate a tufted cell with expression in the soma and primary dendrite extending to the glomerular layer (GL). EPL, external plexiform layer, MCL, mitral cell layer. **b.** Top left: Mean fluorescence image taken in vivo, showing iGluSnFR expression in glomeruli after injection of AAV2.1.hSynap.Flex.SF-iGluSnFR into the OB a Pcdh21-Cre mouse. Top right: ΔF image showing responses to ethyl butyrate (mean of 8 presentations, 2 Hz inhalation). Bottom: Traces showing odorant-evoked iGluSnFR signal in three glomeruli, with dashed region expanded at right. Note distinct temporal responses across successive inhalations for each glomerulus. **c.** Examples of SF-iGluSnFR.A184S responses imaged at high zoom from different glomeruli. Images show mean F (left) and ΔF response maps (right) to ethyl butyrate, showing localization of iGluSnFR signal to the glomerular neuropil (glom.), with a lack of signal on primary dendrites (pr. dend.) or MT cell somata (somas). **d.** Traces showing responses to ethyl butyrate taken from the glomerular neuropil, primary dendrite, and MT somata (locations indicated in (d)), indicating negligible signal on the apical dendrite or somata.

We analyzed inhalation-linked dynamics of the glutamate signal across glomerulus-odor pairs by generating ‘inhalation-triggered average’ responses (ITAs) from multiple inhalations at low frequencies (0.25 - 0.5 Hz)^3,24^. Most ITA transients consisting of a rapid rise and a slower decay to baseline. Odorant-evoked ITA dynamics varied substantially across glomeruli and odorants: glomeruli activated by the same odorant could respond with different onset latencies, rise-times and durations, and the same glomerulus could show distinct ITA dynamics for different odorants (Fig. 2a, b). Inhalation occasionally elicited fluorescence decreases in a glomerulus, suggesting a phasic decrease in ongoing glutamate release (Fig. 2b, see ROI 6), possibly due to the inhalation of clean air into the nasal cavity^24^.

**Fig. 2.**
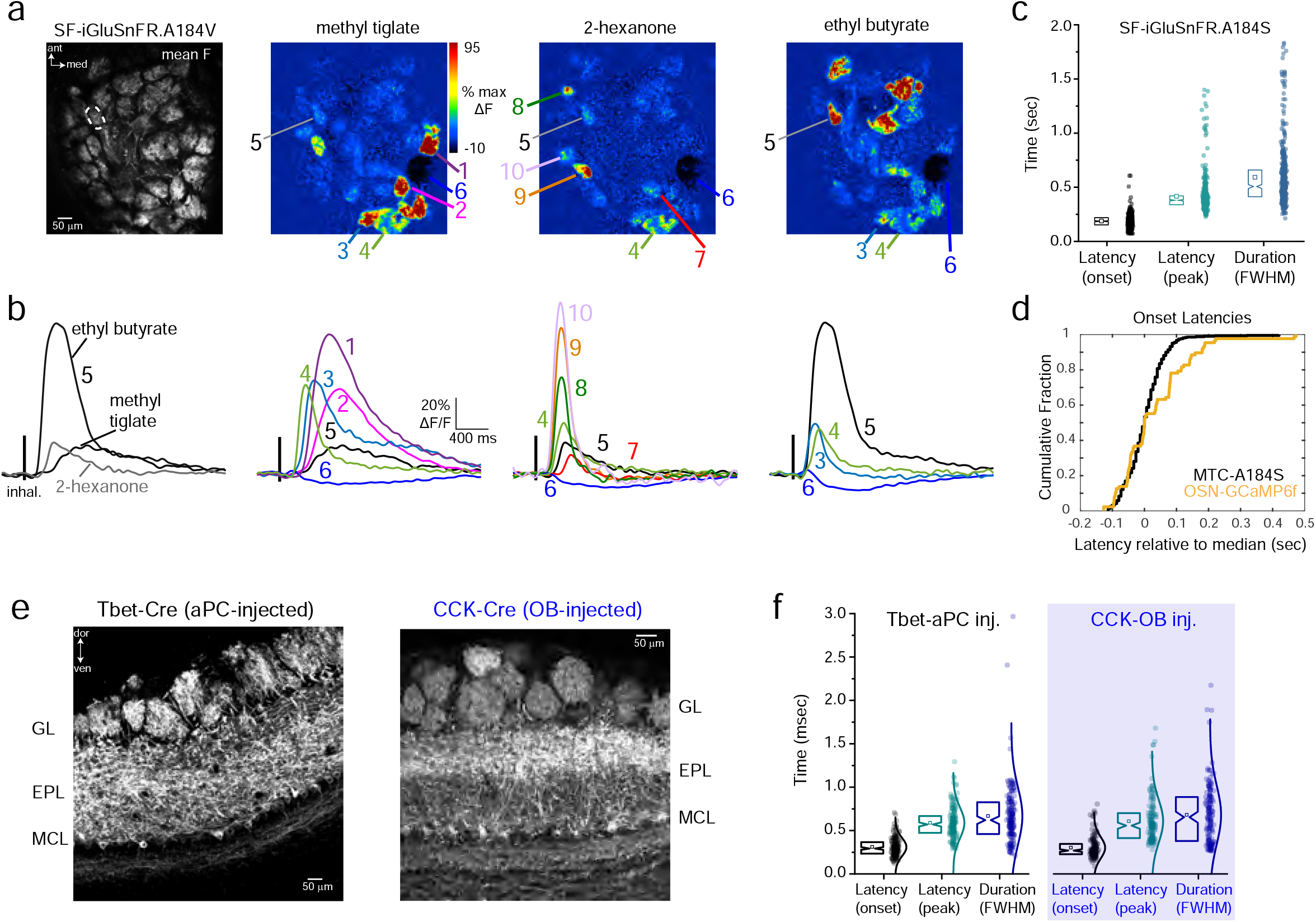
Glomerular glutamate signals show diverse inhalation-driven dynamics. **a.** Left: Mean fluorescence image of SF-iGluSnFR.A184V expression imaged in vivo. Right: Inhalation-triggered ΔF response maps for three odorants. **b.** ITA response traces for the glomeruli indicated in (A). Traces are averages across 17 inhalations. Left: ITAs from one glomerulus (ROI 5) responsive to 3 odorants, illustrating odorant-specific ITA dynamics. Right: ITAs from different glomeruli responsive to each of the three odorants, illustrating glomerulus-specific dynamics. **c.** Plot of ITA onset latency, peak latency, and duration (FWHM, full-width half-max) for all responsive glomerulus-odor pairs using SF-iGluSnFR.A184S. Notch box plots (left) of latency values (right). Notch: median, square: mean, box edges: 25^th^ and 75^th^ percentile, n = 431 glomerulus-odor pairs, 5 mice. **d.** Cumulative fraction of median-subtracted, ITA onset latencies for SF-iGluSnFR.A184S signals, compared to those measured from GCaMP6f expressed in OSN axon terminals (OMP-Cre x Rosa-GCaMP6f mice). Median-subtracted onset latencies were similar: OSNs/GCaMP6f: 0.03 ± 0.11 sec, mean ± SD, n = 91 glomerulus odor pairs, 6 mice; MTs/iGluSnFR.A184S: 0.002 ± 0.06 sec, n = 438 glomerulus-odor pairs, 7 mice, Mann-Whitney Test, p = 0.1. **e.** Preferential iGluSnFR expression in sTCs versus mitral cells. Left: Confocal image of tissue section after Flex.AAV.SF-iGluSnFR injection into the superficial EPL of a CCK-IRES-Cre mouse. Note strongest expression in superficial EPL. Right: Similar image after Flex.AAV.iGluSnFR injection into anterior piriform cortex in a Tbet-Cre mouse. **f.** Identical distributions of onset latencies, times-to-peak, and durations (FWHM) of inhalation-triggered average waveforms imaged from CCK+ sTC and pcMT cell populations. Range of onset latencies (10th - 90th percentiles): 195 – 451 ms (CCK+) vs.192 to 447 ms (pcMTs); times to ITA peak: 373 – 938 ms (CCK+) vs. 367 to 828 ms (pcMTs); FWHM: 313 – 1058 ms (CCK+) vs. 326 to 1007 ms (pcMTs). n = 174 and 160 glomerulus-odor pairs, respectively, from 3 mice each. All statistical comparisons, p>0.05, Kolmogorov-Smirnov test.

We compared ITAs across glomerulus-odor pairs using the three SF-iGluSnFR variants, A184S, A184V and S72A, which have higher, medium and lower affinities for glutamate, respectively^20^. ITA dynamics varied only slightly between the SF-iGluSnFR variants (Supplementary Fig. 1c). We made the most measurements with the medium- and high-affinity variants due to their higher signal-to-noise ratios; across all glomeruli imaged with A184S, ITA onset latencies ranged from 113 to 253 ms (10^th^ - 90^th^ percentiles), time to ITA peak ranged from 300 to 533 ms, and ITA durations ranged from 360 to 867 ms (Fig. 2c). Responses using the A184V variant were similar, with onset latencies ranging from 153 to 313 ms (10^th^ - 90^th^ percentiles, 113 pairs from 3 mice), time to peak ranging from 313 to 727 ms, and ITA durations ranging from 260 to 1100 ms. This overlap suggests that variation in the dynamics of the SF-iGluSnFR signal imaged from MT cell tufts largely reflects the variable dynamics of glutamate signaling in the glomerulus, as opposed to kinetic differences of the reporter. We compared the variability seen in the glutamate ITA with that from OSNs expressing the genetically-encoded reporter GCaMP6f, in response to the same odorants in separate mice (Fig. 2d). Despite the slower kinetics of GCaMP6f compared to iGluSnFR^25^, the variability in onset latencies (relative to median latency) was similar for GCaMP6f and iGluSnFR signals (Fig. 2d). These results suggest that variability in the onset times of glutamate transients onto MT cells after each inhalation largely reflect differences in the timing of spike bursts arriving at OSN presynaptic terminals.

We next asked whether the dynamics of glutamatergic input differ across mitral and tufted cell subpopulations. Mitral cells and superficial tufted cells (sTCs) have distinct odorant response properties^26,27^. Many sTCs express the peptide neurotransmitter cholecystokinin (CCK)^28^; these neurons are strongly driven by monosynaptic input from OSNs^29^, and we have previously shown that they have faster-onset and less diverse odorant-evoked responses than the general MT cell population^6,24,30^. We compared glutamate dynamics onto CCK+ sTCs and piriform cortex-projecting MT cells (pcMTs), using Flex.AAV9.iGluSnFR.A184S virus injection into the superficial EPL of CCK-IRES-Cre mice or retrograde viral expression via injection into anterior piriform cortex of Tbet-Cre mice (Fig. 2e). The latter approach biases expression in mitral and deep tufted cells, with relatively little expression in sTCs^31^. We found no difference in the dynamics of inhalation-linked glutamate transients onto CCK+ sTCs compared to pcMT cells, with both cell types showing a similar distribution of onset latencies, peak times and durations of the iGluSnFR ITA (Fig. 2f). These results indicate that reported differences in odorant-evoked response patterns in sTCs and pcMT cells do not arise from differences in the dynamics of their excitatory input.

### Diversity of glutamatergic signaling across multiple inhalations

To more fully investigate the temporal diversity in patterns of glutamate signaling onto MT cells, we imaged glomerular responses to 23 odorants in single sessions (see Table 1). Odorants were delivered at concentrations such that each odorant evoked relatively sparse responses across the imaging field (Fig. 3a). 16.5% of all glomerulus-odor pairs (748/4520; 7 fields of view, 3 mice) showed responses that were significant according to a conservative criterion of ± 7 SD deviation from baseline. Responsive glomeruli were narrowly tuned across the 23-odorant panel, with high values of lifetime sparseness for excited glomeruli (S = 0.91 ± 0.05, mean ± SD).

**Table 1.**
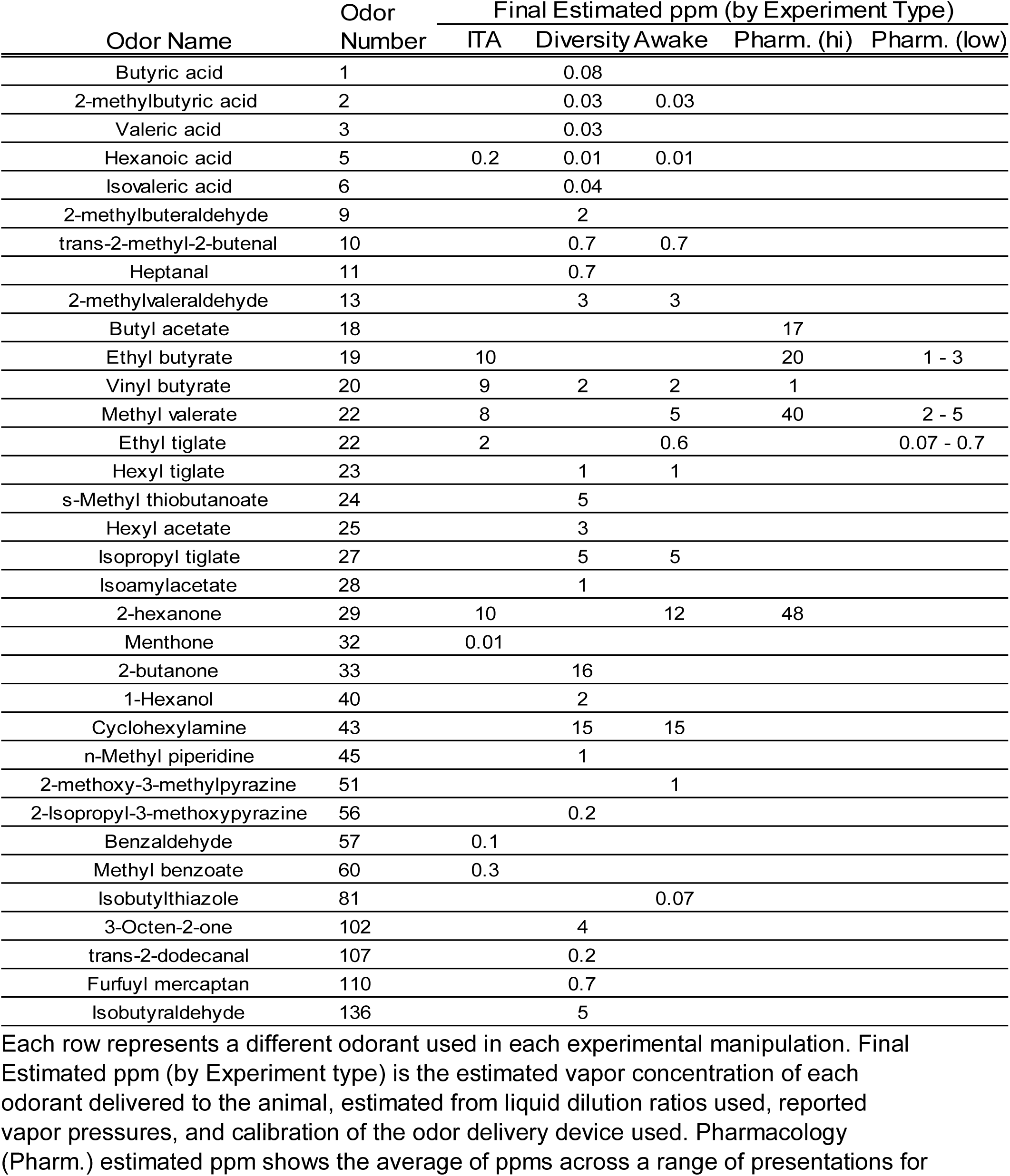
Odorants and concentration used.

**Fig. 3.**
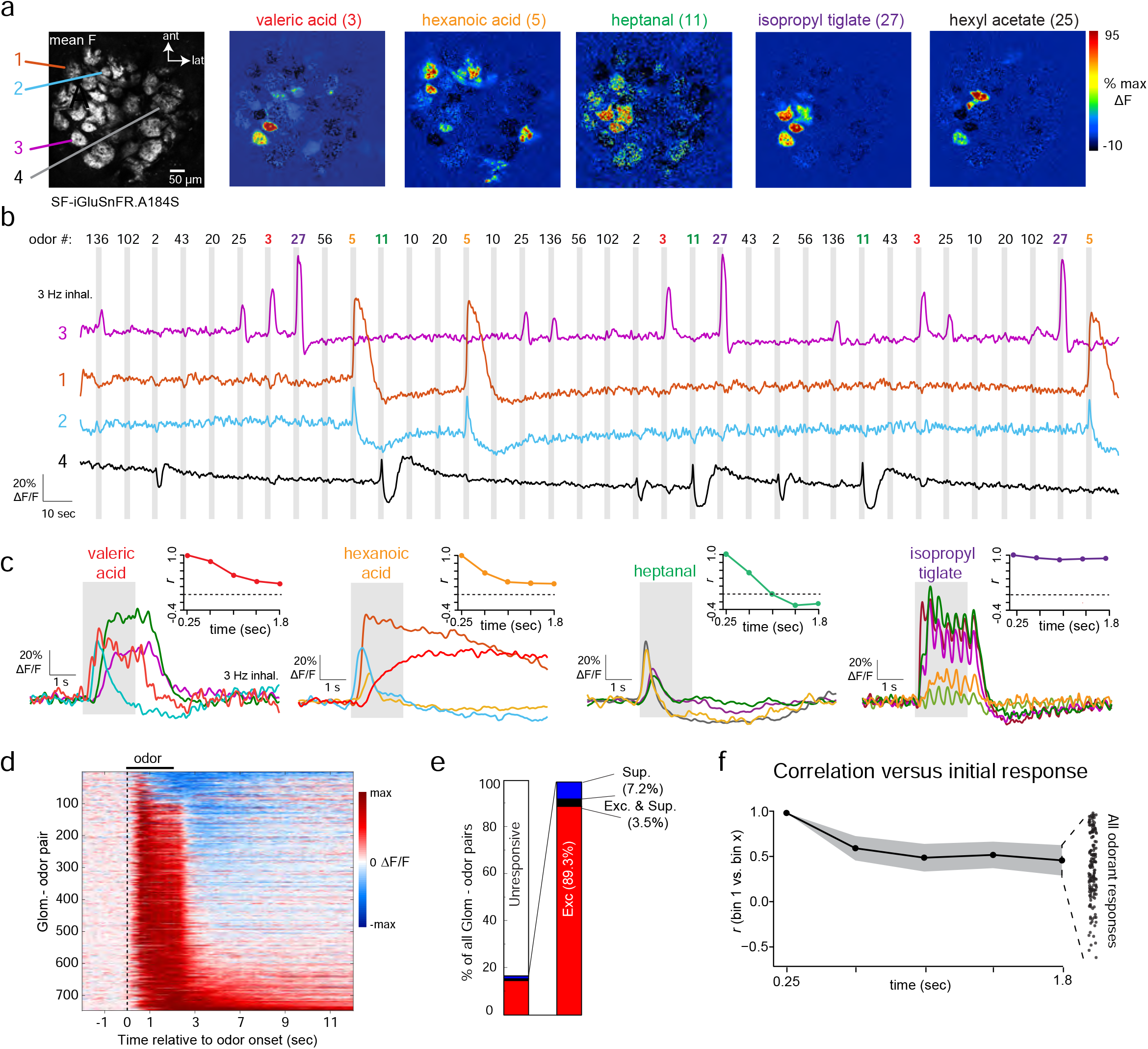
Glomerular glutamate signals show diverse temporal patterns across repeated inhalations of odorant. **a.** Mean fluorescence (left) and ΔF response maps for 5 odorants imaged with SF-iGluSnFR.A184S (right). Odor numbers on the left side of the odorant map indicate odorant identification code for listed odor name (for reference to (B)). Colored text indicates odorants whose responses are shown in panels B, C and F. **b.** Time-series of SF-iGluSnFR.A184S signal from 4 glomeruli, showing continuous signal during 3 repetitions of 12 odorants, presented in random order. Shaded rectangle indicates time of odorant presentations (2 sec duration). Signal is low-pass filtered at 0.5 Hz for display. **c.** Average time-course of odorant-evoked responses in different glomeruli responsive to 4 of the 12 odorants from (B). Insets show change in response pattern across the field of view from (A) over the duration of odorant presentation, expressed as Pearson’s *r* relative to the initial response in successive time bins of 387 ms (see Text). Dotted line indicates r = 0. **d.** ‘Waterfall’ plot showing time-course of odorant-evoked SF-iGluSnFR.A184S signal in all significantly-responding glomerulus-odor pairs (one pair per row), normalized to the peak ΔF/F for each pair. Suppressive responses are low-pass filtered at 0.5 Hz and excitatory responses are low pass filtered at 2 Hz. **e.** Proportion of excitatory (Exc.), suppressive (Sup) and biphasic (Exc. & Sup.) responses across the responsive population (right), as well as the entire population of glomeruli (left). **f.** Change in glutamate response patterns during odorant presentation, summarized over all presentations. Black plot shows mean correlation over time, averaged across all odorant presentations. Shaded area indicates s.e.m. Dot plot at right shows correlation coefficients at the final time bin for all presentations.

Temporal patterns of odorant-evoked glutamate signals were diverse but robust, occurring consistently over repeated presentations (Fig. 3b). The most common response pattern consisted of inhalation-linked transients that persisted over the 2-sec presentation and returned rapidly to baseline (Fig. 3c, e.g., isopropyl tiglate). Other response patterns included transient increases that occurred immediately after odorant onset, prolonged increases that returned slowly to baseline after odorant offset, and slow, ‘facilitating’ rises in glutamate (Fig. 3c, e.g. hexanoic acid). Response patterns varied for different glomeruli activated by the same odorant and for responses of the same glomerulus to different odorants (Fig. 3c), indicating that response patterns were neither a function of SF-iGluSnFR kinetics or expression levels in a particular glomerulus, nor a function of any particular odorant. Odorants also could evoke responses with a pronounced suppressive component (Fig. 3b, c), although these were relatively rare, with pure suppression seen in 7.2% (54/748) of glomerulus-odor pairs and biphasic (excitatory and suppressed) responses seen in 3.5% (26/748) of pairs (Fig. 3d, e).

A consequence of this diversity was that relative glutamate levels in different glomeruli varied over the odor presentation. To quantify this, we generated vectors consisting of time-binned glutamate signals (bin width, 387 ms) across all glomeruli in a field of view and correlated the vector in the first time bin with that from each successive time bin over the 2 sec odor presentation. Among all odorant-fields of view (n = 111), there was a substantial decorrelation beginning in the second bin (time bin 2 vs. time bin 1: Δr = 0.52 ± 0.32) and continuing to the last bin (time bin 5 vs. time bin 1: Δr = 0.39 ± 0.37), although the degree of decorrelation varied greatly for different odorants (Fig. 3f; see also Fig. 3c insets). This decorrelation is qualitatively similar to that observed in patterns of MT cell spiking over repeated inhalations^3,5,6^, suggesting that time-dependent decorrelation can arise at the level of glutamatergic input to MT cells.

### Glomerular glutamate signaling shows diverse temporal dynamics in awake mice

We next imaged iGluSnFR signals in awake, head-fixed mice while monitoring nasal airflow. Respiration frequencies were typically between 3 and 6 Hz (mean frequency per session, 4.4 ± 0.34 Hz, measured from 5 sessions), with occasional pauses and bouts of higher-frequency sniffing. Glutamate transients in awake mice were temporally diverse, similar to anesthetized mice. For many glomeruli, each inhalation at ‘resting’ frequencies (defined as below 5 Hz) elicited a distinct glutamate transient (Fig. 4a, 4b). We measured inhalation-linked dynamics for glomerulus-odorant pairs showing significant respiratory modulation by constructing ITA waveforms as before, but using an external sensor to determine inhalation timing. ITA waveforms generated from these data were approximately sinusoidal (Fig. 4a, lower right), allowing for straightforward estimates of onset latency and time-to-peak of the glutamate signal relative to inhalation. Latency differences between glomeruli were apparent in single trials and were consistent across repeated inhalations of odorant (Fig. 4a). Overall, ITA onset latencies, compiled from four mice, varied over a range of 127 ms relative to the median latency across all glomerulus-odor ITAs in a session (SD = 25 ms, 13 odorants, 7 fields of view, 108 glomerulus-odorant pairs); ITA peak latencies spanned a range of 187 ms (SD = 34 ms) (Fig. 4b). As seen with artificial inhalation, these dynamics could vary across glomeruli and odorants.

**Fig. 4.**
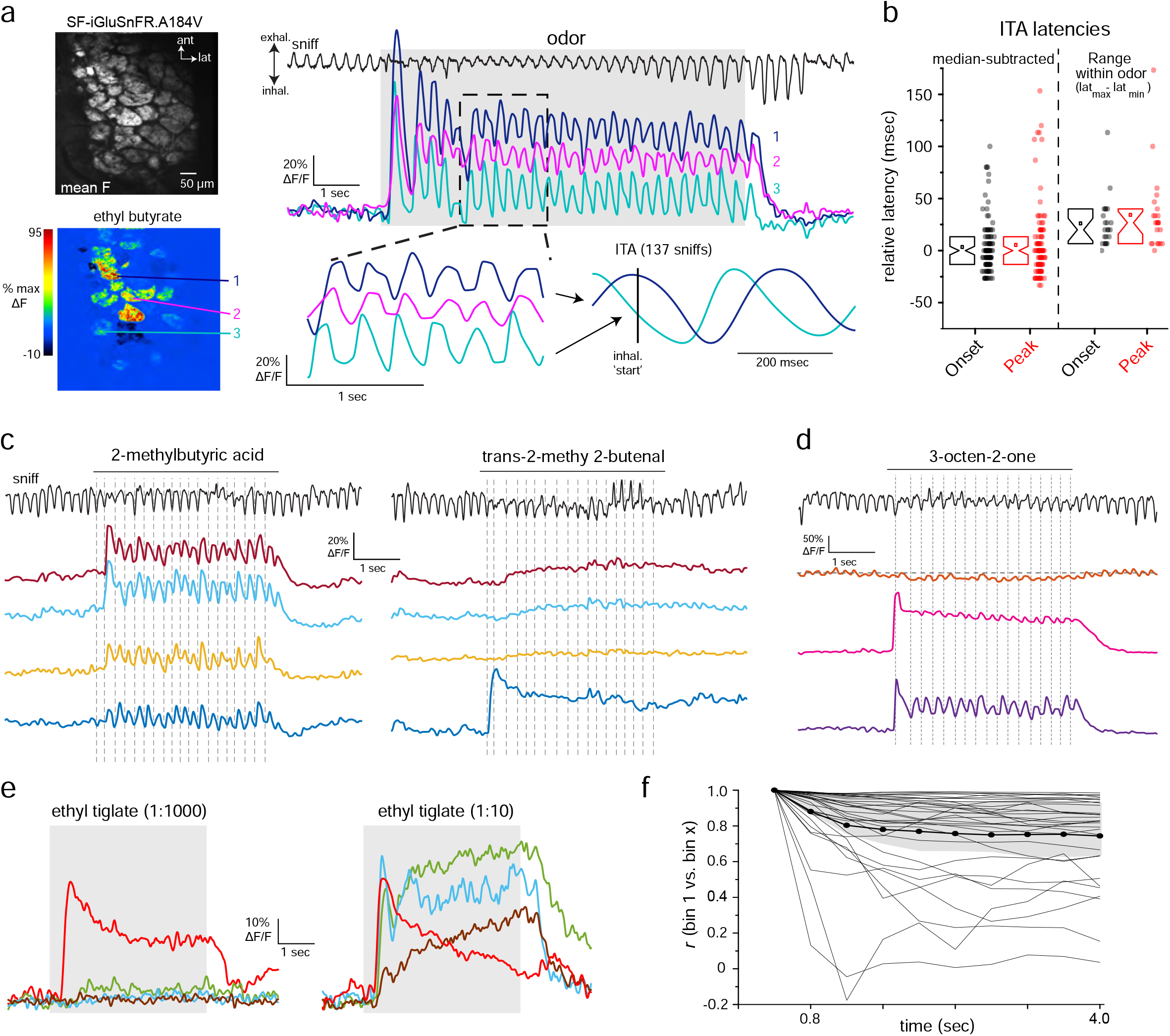
Diversity of glutamate dynamics imaged from MT cells in the awake mouse. **a.** Top: mean SF-iGluSnFR.A184V fluorescence and Bottom: ΔF response map (ethyl butyrate) from the dorsal OB of an awake, head-fixed mouse. Right: Glutamate signals from 3 glomeruli (shown in (A)) during a single presentation of odorant (ethyl butyrate). Top trace shows respiration/sniffing as measured with an external flow sensor, with inhalation oriented downward. Lower left, snippet of signals from each glomerulus, illustrating temporal lag between different glomeruli and consistency of dynamics with each sniff. Signals are unfiltered. Lower right shows ITA waveforms for each glomerulus, generated across 137 sniffs over 6 presentations of ethyl butyrate. Vertical line indicates time of inhalation onset. **b.** Left: Range of ITA onset (black) and peak (red) latencies, relative to median values, compiled across 108 glomerulus-odorant pairs (14 odorants, 7 fields of view). Notch: median, Box edges: 25^th^ and 75^th^ percentiles, square: mean values. Right: Spread of latency values seen across multiple responsive glomeruli imaged in the same field of view for the same odorant (as defined as the difference between maximum (lat_max_) and minimum (lat_min_) latencies), for 23 unique odorant-FOVs. **c.** SF-iGluSnFR.A184V signals imaged from 4 glomeruli from the same FOV showing responses to each of two odorants, showing distinct temporal response patterns across glomeruli and odorants. Traces are unfiltered and from a single presentation. Top trace shows respiration/sniffing for each trial; vertical dotted lines indicate inhalation peak for reference. **d.** SF-iGluSnFR.A184V signals imaged from 3 glomeruli, taken from a separate field of view in the same mouse, showing distinct response patterns to a third odorant (3-Octen-2-one). Note weak suppressive response in one glomerulus (top trace). **e.** Increasing odorant concentration elicits changes in glutamate signal dynamics in awake mice. Traces show averaged responses (8 presentations) from 4 glomeruli in the same field of view to a low (1:1000 dilution) versus high (1:10) concentration of ethyl tiglate. Note that the response in the most sensitive glomerulus shows increased adaptation at the higher concentration, while other glomeruli show facilitating responses. **f.** Changes in glutamate response patterns over the course of odorant presentation, calculated as described for Fig. 3 in 4 awake mice, 33 odorant-FOVs (2 mice, SF-iGluSnFR.A184V; 2 mice, SF-iGluSnFR.A184S). Plots of individual odorant responses are shown in grey, thick black line shows mean, shaded region is quartiles (25^th^ and 75^th^ percentiles). Note high variability in degree of decorrelation over time.

For many glomerulus-odorant pairs, glutamate signals were not modulated by respiration but instead consisted of a tonic glutamate increase. The same glomerulus could show strong coupling to respiration for one odorant and a lack of respiratory coupling for another odorant (Fig. 4c). Similarly, different glomeruli responsive to the same odorant could show either strong or no respiratory coupling (Fig. 4d, indicating that the lack of coupling was not due to differences in respiration from trial to trial, nor was it due to differences in odorant stimulus profile. Odorants could also elicit decreases in the glutamate signal (Fig. 4d, top trace). Respiration-coupled and tonic glutamate signals were apparent to a similar degree in mice expressing SF-iGluSnFR.A184V or SF-iGluSnFR.A184S (2 mice each).

Glutamate signals in awake mice also showed diverse odorant responses over multiple inhalations, including slowly facilitating and adapting response patterns (Fig. 4c, d). We also observed concentration-dependent changes in glutamate dynamics, with increasing odorant concentrations leading to stronger adaptation of glutamate signal for strongly-responsive glomeruli and recruitment of slowly-increasing responses in weakly-responsive glomeruli (Fig. 4e). Consequently, the relative patterns of glutamate signals across glomeruli changed from their initial pattern over repeated sniffs, and glutamate response patterns decorrelated more substantially for some odorants than others (Fig. 4f). Overall, these data indicate that the diversity in dynamics of glutamatergic inputs to MT cells observed in awake mice is qualitatively similar to that imaged with artificial inhalation.

### Correspondence between dynamics of glutamatergic inputs and MT cell postsynaptic activity

While glutamatergic signaling in the glomerulus is the sole source of excitatory input to MT cells, inhibitory circuits and MT cell intrinsic properties may also shape patterns of MT cell excitation^2,10,11,32,33^. To assess these relative contributions, we compared glutamate signals with calcium signals measured from MT cell apical tufts of the same glomerulus, imaged simultaneously. Because MT cell spikes back-propagate along the MT cell primary dendrite and invade the apical tuft, calcium signals imaged from the tuft provide a proxy for patterns of MT cell spike output from each glomerulus^34-36^.

We co-expressed SF-iGluSnFR.A184S and the red calcium reporter jRGECO1a^37^ in MT cells using viral co-injection into the OB of Tbet-Cre mice (Fig. 5a). In response to odorant stimulation, jRGECO1a signals were apparent both in an activated glomerulus as well as in the primary dendrites of the MT cells innervating it, while SF-iGluSnFR signals were not apparent outside the glomerular neuropile (Fig. 5b, e). As in our earlier report^30^, we saw very high correspondence between odorant-evoked GCaMP6f signals imaged from the soma or primary dendrite of MT cells and those imaged from the neuropil of their parent glomerulus (Fig. 5b; Supplementary Fig. 3). In anesthetized mice, inhalation-linked transients were apparent in both signals at 1 Hz inhalation, although the jRGECO1a calcium signal decayed more slowly (Fig.5b). Despite these kinetic differences, there was a close correspondence in the inhalation-linked dynamics of the two signals, with different glomeruli showing the same relative differences in the onset latency and time to peak of the SF-iGluSnFR and jRGECO1a ITA waveform (Fig. 5c, d). We also observed a close - though not perfect - correspondence in the occurrence of spontaneous transients in the SF-iGluSnFR and jRGECO1a signals (Supplementary Fig. 2a). In awake mice, jRGECO1a signals showed little or no inhalation-linked modulation despite strong inhalation-linked transients in the SF-iGluSnFR signal (Fig. 5f).

**Fig. 5.**
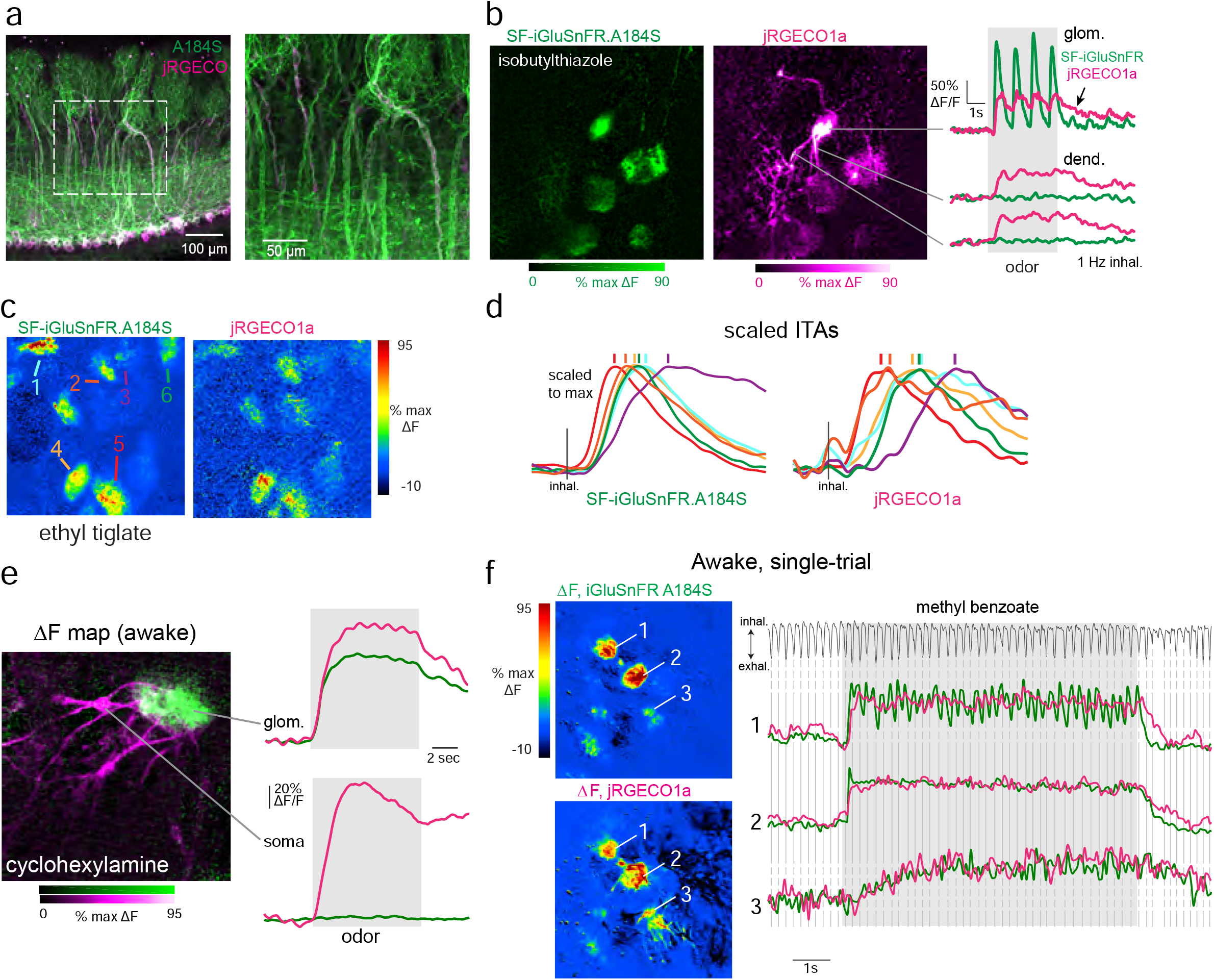
Dual-color imaging reveals high correspondence between pre- and postsynaptic signals in MT cells of the same glomerulus. **a.** Post-hoc confocal image showing coexpression of jRGECO1a (magenta) and SF-iGluSnFR.A184S (A184S, green) in MT cells of a Tbet-Cre mouse. **b.** Dual-color two-photon imaging of SF.iGluSnFR.A184S and jRGECO1a signals from MT cells. Left: odorant-evoked ΔF response maps for SF-iGluSnFR and jRGECO1a signals (magenta imaged simultaneously. Note jRGECO1a signal in dendrites of several MT cells exiting the central glomerulus, with SF-iGluSnFR signal confined to the glomerular neuropil. Right: traces showing time-course of the fluorescence signal in each channel (1 Hz inhalation). Top traces are from glomerular neuropil; lower traces are from dendrites outside of the glomerulus. **c.** Pseudocolor odorant-evoked ITA response maps across the green (A184S) and red (jRGECO1a) channels. Arrows indicate ROIs with traces plotted in (d). **d.** ITA traces taken from different glomeruli activated by the odorant in (c), with different onset latencies, times to peak and durations in different glomeruli. Left traces: SF-iGluSnFR.A184S; right traces: jRGECO1a; traces from the same glomerulus are shown with the same color in each set. Each ITA trace is scaled to the same maximum. Vertical lines indicate peak time for the signal in each glomerulus. The relative order of peak times is the same for both signals. **e.** Dual-color imaging of SF.iGluSnFR.A184S and jRGECO1a signals from MT cells in the awake mouse. Left: Composite dual-color ΔF response map showing SF-iGluSnFR (green) and jRGECO1a (magenta) signals evoked by cyclohexylamine. Right: Traces showing fluorescence signal taken from the glomerular neuropile (top) and soma (bottom) of an innervating TC. Traces are average of 8 presentations. **f.** Trial-averaged ΔF response maps and traces for SF-iGluSnFR.A184S and jRGECO1a signals imaged from a single presentation of odorant in an awake mouse. Top trace shows respiration measured via external flow sensor. The SF-iGluSnFR signal clearly follows each inhalation in only one of the three glomeruli shown, while the jRGECO1a signal does not follow individual inhalations well in any glomeruli.

To compare odorant-evoked response patterns over a time-scale involving multiple inhalations, we first performed dual-color imaging in anesthetized mice using the same 23 odorant panel as for the single-color imaging (5 fields of view from 3 mice). Overall, there was a striking degree of correspondence in response patterns of the two signals, with only rare exceptions (Fig. 6a; Supplementary Fig. 2b). There was high concordance in glomeruli showing purely excitatory responses as measured with SF-iGluSnFR and jRGECO1a (using the conservative criterion of ± 7 SD above baseline): 70% (273/387) of glomerulus-odor pairs with an excitatory SF-iGluSnFR signal also showed a significant jRGECO1a response, and 79% (273/346) of glomeruli with significant jRGECO1a increases showed excitatory SF-iGluSnFR signals (Supplementary Fig. 2c). In awake mice, using a smaller odorant panel, only 46% (57/123) of glomerulus-odor pairs with a significant excitatory SF-iGluSnFR signal also showed jRGECO1a responses; however, 93% (57/61) of glomerulus-odor pairs with excitatory jRGECO1a responses also showed SF-iGluSnFR responses. Temporal response patterns were also highly correlated in awake mice (Fig. 6b; Supplementary Fig. 2d); we quantified this using an index, T_2_-T_1_/T_max_, that distinguished sustained, rapidly adapting, and facilitating responses. T_2_-T_1_/T_max_ values were highly correlated for glutamate and Ca^2+^ signals measured for the same glomerulus-odor pair, both in anesthetized and awake mice (Fig. 6c). These results suggest that the evolution of MT cell activity patterns across multiple inhalations largely follows that of glutamatergic input to these cells.

**Fig. 6.**
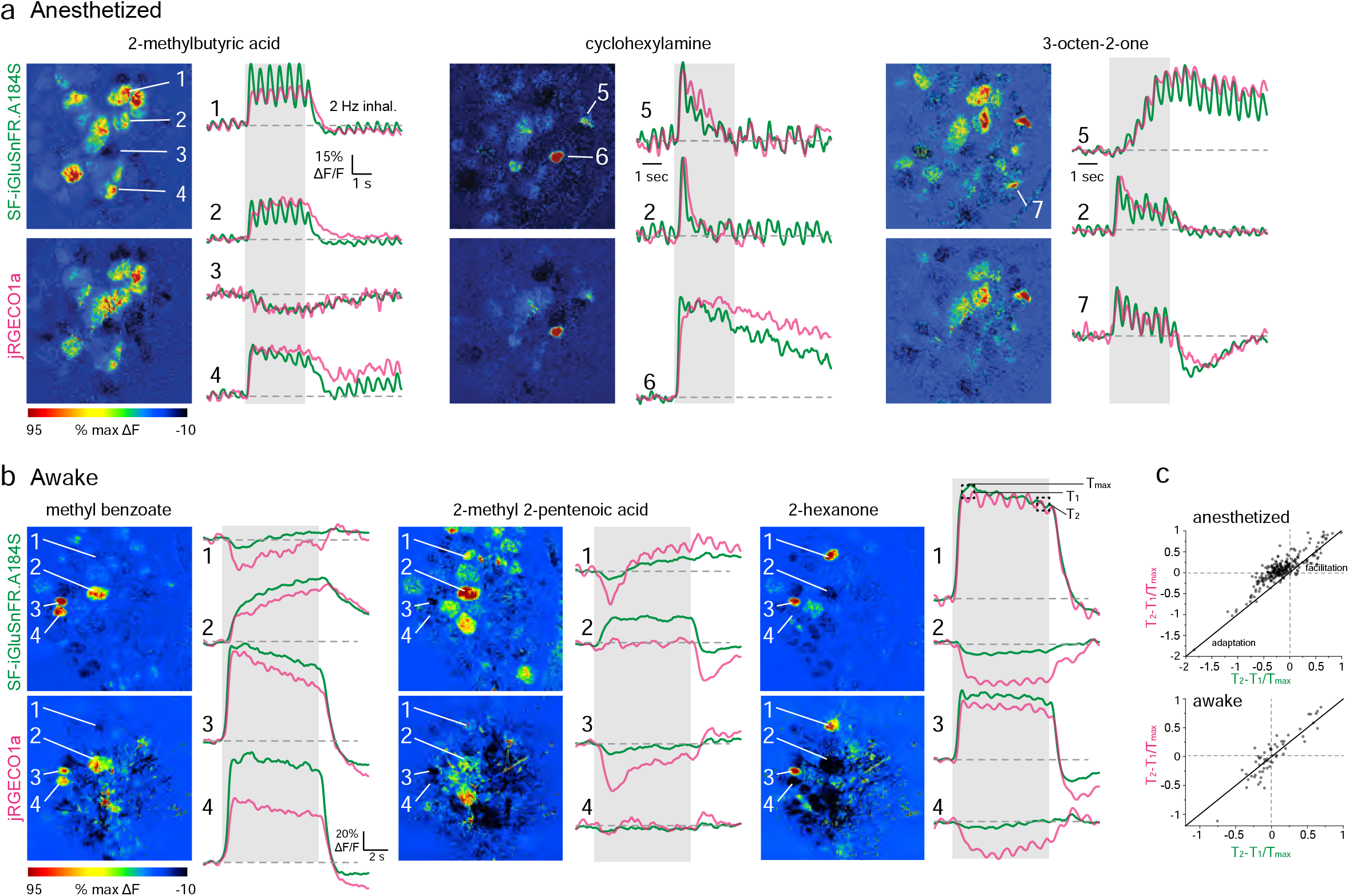
High correspondence between odorant-evoked temporal patterns of glutamate signaling and calcium activity in MT cells of the same glomerulus. **a.** ΔF response maps (left) and trial-averaged SF-iGluSnFR.A184S and jRGECO1a signals imaged from select glomeruli for three odorants, illustrating high correspondence in slow response dynamics. Traces are average of 4 presentations in the same anesthetized mouse. Numbers indicate different glomeruli. Traces for cyclohexylamine and 3-octen-2-one are scaled to the same maximum. **b.** ΔF response maps and trial-averaged SF-iGluSnFR.A184S and jRGECO1a responses taken from four glomeruli in response to three odorants in an awake mouse. Traces are average of 16 presentations. Note distinct response patterns for the same glomerulus in response to different odorants, but, with a few exceptions, similar response patterns for SF-iGluSnFR and jRGECO1a signals. **c.** High correlation between T2-T1/T_max_ values measured for excitatory SF-iGluSnFR and jRGECO1a responses in the same glomerulus in anesthetized (top) or awake mice (bottom). Correlation coefficients for T2-T1/T_max_ values between SF-iGluSnFR and jRGECO1a responses were r = 0.87 in anesthetized mice (273 pairs,3 mice) and r = 0.90 in awake mice (57 pairs, 2 mice).

To infer how inhibitory circuits might shape MT cell spike output from glomeruli, we compared the prevalence of suppressive response components in the glutamate and Ca^2+^ signals. In anesthetized mice, suppressive responses were rare, and their prevalence may be underestimated from the relatively few repeat trials given (3 - 4 per odorant) and the strict significance criteria we used. Nonetheless, jRGECO1a responses with suppressive components were detected significantly more frequently than SF-iGluSnFR responses, being present in 1.8% (72/3840) versus 0.7% (27/3840) of all glomerulus-odor pairs, and 17% versus 7% of all significant responses (Chi-Square statistic = 20.45, p = 6.0 e^-6^). In awake mice, suppressive responses were more prevalent (Supplementary Fig. 3d), with a similar prevalence of suppressive components in both signals, being present in 21% (jRGECO1a: 92/446) versus 17% (SF-iGluSnFR: 78/446) of all glomerulus-odor pairs (Chi-Square statistic = 1.15, p = 0.28). Notably, suppressive jRGECO1a responses were very rarely seen in glomeruli showing excitatory SF-iGluSnFR signals, detected in only 2 of 72 suppressive jRGECO1a responses in anesthetized mice (2.7%) and 5 of 92 responses (5.4%) in awake mice; this prevalence is approaching the false positive rate of our measure of response significance. This result is surprising given predictions from slice experiments that weak excitatory input can lead to MT cell suppression by driving feedforward inhibition^11,17,38^. However, these results are consistent with a model in which interglomerular inhibition suppresses MT cell spiking in a small fraction of glomeruli^30,39,40^.

### Limited contribution of multisynaptic pathways to odorant-evoked glutamate signaling onto MT cells

Sensory-evoked glutamatergic input onto MT cells can arise from OSNs or from multisynaptic excitation involving dendritic glutamate release from ET cells and, possibly, sTCs^16,29,41-43^. In addition, inhibitory circuits can modulate multisynaptic glutamate signaling^11,12,17^. Glomerular glutamate signals may also arise from glutamate release by MT cell dendrites^44,45^, and thus at least partially reflect MT cell excitation itself. To isolate the direct contribution of OSN inputs to MT cell excitation dynamics, we compared evoked glutamate dynamics before and after pharmacological blockade of postsynaptic activity with ionotropic glutamate receptor antagonists^46-48^. Because iGluSnFRs are insensitive to these antagonists^19,20^, we reasoned that this approach would largely prevent OSN-driven excitation of postsynaptic neurons and allow us to image evoked MT cell glutamate signals arising solely from OSN inputs.

To confirm postsynaptic blockade, we tested the ability of APV+NBQX to block activation of CCK-expressing sTCs, as these, like ET cells, are strongly driven by monosynaptic OSN input^29^. We imaged activation of CCK+ OB neurons in CCK-IRES-Cre mice crossed to a Cre-dependent GCaMP6f reporter line. We first applied the GABA_B_ receptor antagonist CGP35348 (1 mM) to the dorsal OB to further enhance transmitter release from OSNs by removing presynaptic inhibition, as done previously^23,49^. As expected, CGP35348 led to a modest (20%) increase in the peak CCK+ GCaMP6f response (Fig. 7b). Subsequent application of APV+NBQX (1 mM/0.5 mM) had mixed effects in different glomeruli, with responses almost completely eliminated in 8 of 14 glomeruli (96% mean reduction in peak response, 3 experiments; see Fig. 7c legend for summary statistics) but only partially reduced in the remaining 6 (46% mean reduction) (Fig. 7b - d). We performed additional experiments using a 5x higher concentration of antagonist (5 mM APV/2.5 mM NBQX) and without preapplication of CGP35348. In these, APV+NBQX completely blocked postsynaptic activation in 11 of 13 glomeruli (from 3 experiments), with strongly reduced responses in the remaining two (Fig. 7e).

**Fig. 7.**
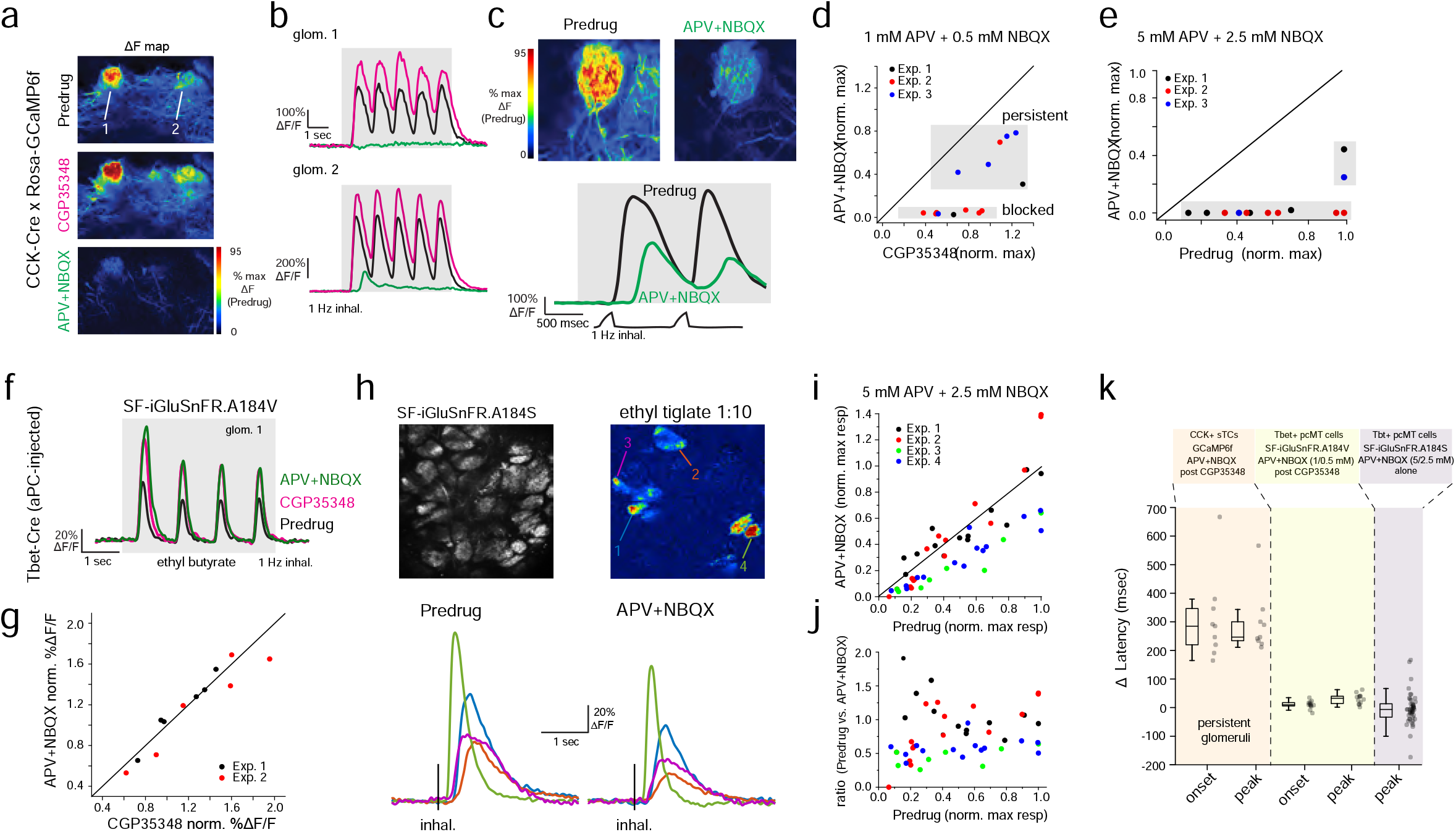
Effects of pharmacological blockade of polysynaptic glutamate signaling onto MT cells. **a.** Odorant response maps of GCaMP6f signals imaged from CCK+ sTCs, taken across predrug, CGP35348 and APV+NBQX conditions, scaled to predrug response levels. **b.** Response traces for the two glomeruli indicated in (k), showing enhanced response amplitudes after application of CGP35348 (magenta) and either elimination (top) or strong reduction (bottom) in responses after subsequent application of APV+NBQX (green). In the latter case, the remaining response after APV+NBQX is delayed. **c.** Example from a second experiment of a glomerulus with persistent responses after APV+NBQX application. Traces (below) show expansion of response to first two inhalations of odorant, showing delayed GCaMP6f response after APV+NBQX. **d.** Scatter plot comparing CCK+ GCaMP6f responses (peak amplitude, first inhalation) across CGP35348 and subsequent APV+NBQX conditions, imaged from 14 glomeruli in 3 experiments (2 mice). Summary statistics: ratio of post:pre APV+NBQX amplitudes, 0.27 ± 0.27, Wilcoxon signed ranks Z = 3.26, p = 0.001, n = 14 glomeruli). Shaded regions indicate two glomeruli with near-totally blocked responses or with persistent but weakened responses after APV+NBQX. Median post:pre APV+NBQX ratios, persistent: 0.54 ± 0.16 (n = 6), blocked: 0.06 ± 0.03 (n = 8). **e.** Scatter plot of CCK+ GCaMP6f responses before and after APV+NBQX (5 mM/2.5 mM), without prior application of CGP35348. **f.** Traces showing iGluSnFR.A184V signals imaged from pcMT cells in an aPC-injected Tbet-Cre mouse, comparing responses to the same odorant across Predrug, CGP35348, and APV+NBQX conditions. Inhalation frequency, 1 Hz. **g.** Comparison of peak iGluSnFR responses in pcMT cells across CGP35348 and APV+NBQX conditions for first inhalation of odorant. Points indicate glomeruli from two mice (red and black). APV+NBQX/CGP35348 ratios, mouse 1 median: 1.03 (range, 0.89 - 1.11, n = 6), mouse 2: 0.86 (range, 0.79 - 1.05, n = 6). Combined comparison: Wilcoxon signed ranks, Z = 0.59, p = 0.55, n = 12. CGP35348/predrug ratios for the two mice (not shown), mouse 1 median: 1.32 (range, 1.28 - 1.52), mouse 2 median: 1.80 (range, 1.59 - 2.33); Combined comparison: Wilcoxon signed ranks, Z = -3.02, p = 0.003, n =12. **h.** Top: Mean fluorescence and ΔF response maps showing odorant-evoked iGluSnFR signals in several glomeruli after SF-iGluSnFR.A184S expression in piriform-projecting MT cells. Bottom: Inhalation-triggered average responses from four of the glomeruli, imaged before and after application of APV+NBQX (5 mM / 2.5 mM). Response amplitudes in each glomerulus are modestly reduced, but relative latencies and durations are unchanged. **i.** ITA response amplitudes before and after APV+NBQX (5 mM / 2.5 mM) application in pcMT cells from four experiments (47 total glomeruli). In each experiment, the same odorant was presented at two concentrations varying by a factor of 2.5 – 10; responses are plotted normalized to the maximal pre-drug response in each experiment. Summary statistics, paired Wilcoxon signed ranks per experiment, Exp. 1: n = 13, z = 0.21, p = 0.83; Exp. 2: n=16, z=0.18, p=0.86; Exp. 3: n = 8, z=2.5, p=0.008; Exp. 4: n = 14, z=3.3, p=1.2e-4. **j.** Same data as in (i), showing ratio of post- to pre-drug ITA amplitude as a function of pre-drug response. Scatter indicates lack of consistent effect of APV + NBQX as a function of initial response amplitude. **k.** Comparison of change in onset latencies and peak times of inhalation-linked responses after APV+NBQX application, measured from CCK+ GCaMP6f datasets (persistent glomeruli, (**d, e**)), pcMT iGluSnFR with 1 mM/0.5 mM APV+NBQX (data in (**g**)), and pcMT iGluSnFR with 5 mM/2.5 mM APV+NBQX (data in (**i**)). Note no change in latencies for iGluSnFR signals. Boxes indicate 25^th^ and 75^th^ percentiles, line indicates median, and whiskers denote outliers with a coefficient of 1.5. Summary statistics: 1) CCK+ GCaMP6f, APV+NBQX vs. CGP35348, median Δ onset latency: 285 ms (range, 165 – 667 ms, n = 9 glomeruli), Z = -2.6 p = 0.004, Δ peak latency: 247 ms (range, 211 − 567 ms), Z = -2.6, p = 0.004. 2) iGluSnFR pcMTs, APV+NBQX vs. CGP35348 Δ onset latency: mouse 1: 9.9 ms (range, -9.1 – 34.7 ms, n = 6), mouse 2: 9.8 ms (range, -19.4 − 18.6 ms, n = 6); Z = -1.47 p = 0.14 (same for both mice); Δ peak latency: mouse 1: 40.05 ms (range, 25.4 – 62.6 ms, n = 6), mouse 2: 14.5 ms (range, 1.8 – 38.8 ms, n = 6); Z = -1.89, -2.10, p = 0.06, 0.04); 3) iGluSnFR pcMTs, pre- vs. post- APV+NBQX, median Δ peak latency, -6.7 ms; paired Wilcoxon signed ranks: expt. 1, n = 13, z=1.64, p=0.1; expt. 2: n = 14, z=-0.55, p=0.58; expt. 3: n = 6, z=00, p=1; expt. 4: n = 14, z=1.09, p=0.27.

Importantly, in glomeruli with persistent (but weakened) responses after APV+NBQX application, inhalation-linked glutamate transients were significantly delayed compared to the predrug condition (Fig. 7c, k), with a median increase in onset latency of 285 ms (range: 164 – 667 ms) and median increase in time to peak of 247 ms (range: 211 – 567 ms). An explanation for this substantial delay is that, in glomeruli receiving the strongest sensory input, glutamate concentrations are eventually able to overcome APV+NBQX blockade sufficiently to trigger spike bursts in sTCs. In additional experiments imaging GCaMP6f signals from the general population of Tbet-expressing MT cells, APV+NBQX eliminated inhalation-linked transients in glomeruli, with only a small-amplitude tonic glutamate signal remaining (Supplementary Fig. 4 a,b), possibly mediated by metabotropic glutamate receptors. These results demonstrate that APV+NBQX blocks postsynaptic activation in glomeruli receiving all but the strongest OSN inputs, and even in those glomeruli, substantially weakens and delays activation of monosynaptically-driven TCs.

We next used APV+NBQX blockade to test the contribution of mono-versus di- or polysynaptic glutamate signaling to inhalation-linked glutamate transients onto pcMT cells. We first tested effects of CGP35348 and subsequent application of APV+NBQX (1 mM/0.5 mM) on glutamate transients using the medium-affinity SF-iGluSnFR.A184V. Inhalation-linked glutamate transients were increased by CGP35348, confirming earlier results^23^. Surprisingly, subsequent application of APV+NBQX had no impact on response amplitudes (Fig. 7f, g). Importantly, GABA_B_ receptor or iGluR blockade had little to no effect on inhalation-linked glutamate dynamics (Fig. 7f, k). We obtained similar results in epifluorescence experiments imaging iGluSnFR signals from a Tbet-expressing MT cell population across a larger area of the dorsal OB, using epifluorescence imaging (Supplementary Fig. 4d-f).

Multisynaptic glutamatergic excitation may preferentially drive MT cell responses to weak inputs^17,43,50^, and the odorant concentrations used in the preceding experiments were relatively high (20 – 50 ppm see Table 1). Removal of presynaptic inhibition with CGP35348 may also enhance OSN input strength to unnaturally high levels. Thus, we next tested the impact of glutamate receptor blockade on inhalation-linked transients in pcMT cell responses to odorants presented at 10 – 100x lower concentrations (0.1 – 5 ppm), using a higher concentration of APV+NBQX (5 mM/2.5 mM) and omitting CGP35348. Here, iGluR blockade was weakly effective at reducing the magnitude of inhalation-linked glutamate transients, causing a significant reduction in two of four experiments and a mean reduction of 32 ± 12% overall (mean ± s.e.m. of median ratios across 4 experiments, n = 8 – 16 glomerulus-odor pairs per experiment) (Fig. 7i, j). However, APV+NBQX had no impact on the dynamics of inhalation-linked glutamate responses, with no change in peak times in any experiment (Fig. 7k).

Finally, to examine the role of direct versus multisynaptic excitation in shaping diverse temporal patterns of glutamatergic input to MT cells, we used a 12-odorant panel (delivered concentrations ranging from 0.03 - 15 ppm, see Table 1) and 3 Hz artificial inhalation, comparing response patterns before and after application of APV+NBQX (1 mM/0.5 mM). This approach yielded significant odorant responses in 83 glomerulus-odorant pairs across 3 mice, and included diverse excitatory response patterns (Fig. 8a). Consistent with the earlier experiments, APV+NBQX had little overall impact on the magnitude of evoked iGluSnFR signals: while there was some variability in the effect for particular glomerulus-odor pairs, neither mean nor peak excitatory response magnitudes showed a significant overall change after APV+NBQX application (paired t-tests, p = 0.147 for mean amplitude, p = 0.605 for peak amplitude, n = 83) (Fig. 8b).

**Fig. 8.**
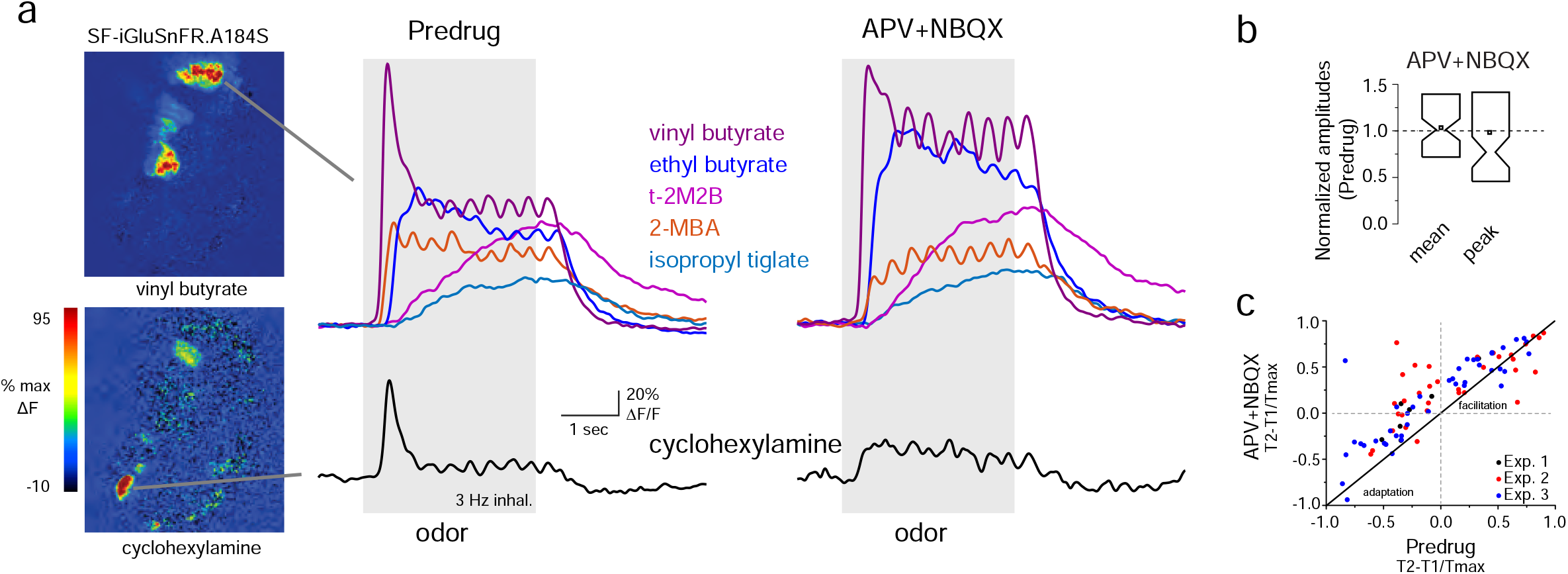
Polysynaptic glutamate signaling contributes to responses to low-concentration odorants and to slow temporal dynamics across inhalations. **a.** Left: ΔF response maps showing responses to two odorants before application of APV+NBQX (1 mM/0.5 mM). Signals are from SF-iGluSnFR.A184S injected into the OB of a Tbet-Cre mouse. Right: Traces (mean of 4 presentations) from two glomeruli (top, bottom) showing diverse temporal responses to different odorants, before and after APV+NBQX application. Response to cyclohexylamine (bottom) shows a loss of the initial transient and replacement by tonic signal. Response to vinyl butyrate (top) shows increase in initial response magnitude and apparent loss of adaptation. Temporally distinct patterns of response patterns to other odorants persist after drug application. **b.** Mean and peak normalized response amplitudes after application of APV+NBQX, with distributions indicated as previously. **c.** Scatter plot of T_2_-T_1_/T_max_ values before and after APV+NBQX for all responsive glomerulus-odor pairs across 3 mice (each mouse indicated by color), showing decrease in adaptation for many pairs after application of APV+NBQX. Summary statistics: paired t-tests, mouse 1, n= 7 glomerulus-odor pairs, p=0.001; mouse 2: n = 34, p=0.001; mouse 3: n = 42, p=5.45 x 10^−6^.

APV+NBQX did, however, impact the temporal patterns of the glutamate signal for a subset of responses. In particular, APV+NBQX tended to reduce adaptation that occurred following the initial inhalation of odorant or enhance longer-lasting, tonic-type responses (Fig. 8a). This was reflected as an increase in T_2_-T_1_/T_max_ values by APV+NBQX, with significant increases seen in each of three mice (Fig. 8c). Taken together, these results suggest that multisynaptic glutamatergic signaling contributes rather subtly to odorant-evoked excitation of MT cells, slightly enhancing weak inhalation-linked glutamate transients but also contributing to adaptation of glutamate signals during repeated odorant sampling.

## DISCUSSION

Temporally complex patterns of excitation and inhibition among principal cells (MT cells) of the OB are hypothesized to play important roles in coding olfactory information, and have been attributed to emergent properties of excitatory and inhibitory circuit interactions^4-6,30,33^. By directly imaging glutamate signaling onto MT cell dendrites of OB glomeruli, we observed a striking degree of heterogeneity in this signaling both on the time-scale of a single respiratory cycle and over a slower time scale involving repeated samples of odorant. Simultaneous imaging of presynaptic glutamate and postsynaptic calcium from MT cells in the same glomerulus showed a high correspondence in the dynamics of glutamatergic input MT output. Pharmacological blockade of postsynaptic activity had little impact on glutamate dynamics, implicating direct inputs from OSNs as a primary source of this diversity. These results suggest a model of OB circuit function in which the dynamics of sensory input to glomeruli can account for much of the diversity of MT cell responses that underlies odor representations in vivo.

The iGluSnFR sensors are well-suited to characterizing glutamatergic signaling in the OB in vivo, with intrinsic kinetics (rise and decay times of <10 to ∼ 20 ms, respectively^19,51,52^) that are much faster than the dynamics of odorant-evoked excitatory input to MT cells (EPSP rise- and decay times of ∼100 ms and 200 ms, respectively^53^). iGluSnFR expression can prolong the decay of glutamate transients by competing with glutamate transporters^54^, and decay rates are slower (∼140 ms) for the high-affinity SF-iGluSnFR.A184S ^20,55^. While we did find that decay times skewed longer for ITA responses measured with the high-affinity variant, their distribution largely overlapped with that of the medium-affinity A184V. Furthermore, the degree of respiratory modulation as well as overall diversity in slow iGluSnFR signal dynamics appeared similar for the two variants. Thus, the SF-iGluSnFR signal imaged from MT cell apical tufts is a reliable reporter of the variable dynamics of glutamate signaling in the glomerulus.

Another potential confound to interpreting the iGluSnFR signal is dendritic release of glutamate from MT cells themselves^44,45^. However, near-complete pharmacological blockade of MT cell activation did not alter ITA waveforms, suggesting that glutamate signals imaged from OB glomeruli reflect glutamatergic input to, as opposed to glutamate release from, MT cells.

OSN inputs to the OB respond with different latencies relative to inhalation in a glomerulus- and odorant-specific manner^15,24,56^, and we found a close match in the range of response latencies for SF-iGluSnFR signals and those measured from OSN presynaptic terminals using Ca^2+^ reporters. We also saw variation in other aspects of inhalation-linked glutamate dynamics that are not predicted from presynaptic Ca^2+^ imaging, with the duration of inhalation-linked glutamate transients ranging from 200 ms to over 1.5 sec. Glutamatergic signaling onto MT cells also showed complex dynamics over a slower time scale spanning multiple inhalations that could include adaptation, facilitation, or biphasic response patterns. This diversity likely has substantial impacts on the temporal dynamics of MT cell spiking, and suggests that MT cell excitation is not a simple linear combination of responses to a single inhalation^57^.

Several results emerged from imaging glutamate dynamics in awake mice that inform models of how olfactory information is represented during natural odor sampling. First, the relative magnitudes of glutamate signal across different glomeruli changed over the course of an odor presentation, suggesting that odorant representations evolve rapidly during repeated sampling of odorant. This result is consistent with recent findings of dynamic odor representations on a similar time scale at the level of MT cell spiking^3,5^. Second, we found that a substantial fraction of odorant-evoked glutamate signals showed minimal or no respiratory patterning, instead showing tonic increases in glutamate when odorant was present. Assuming a similar fraction of MT cells fail to show respiratory patterning in their spike timing in awake mice, this result has implications for theories of odor identity or intensity coding that are based on the timing of MT cell spikes within the breathing cycle^4,7-9^. Third, in both awake and in anesthetized mice, we found that the temporal evolution of glutamate responses as well as the degree of inhalation-linked patterning varies with odorant identity and concentration as well as glomerular identity. Comparable richness in response kinetics have been well-characterized in Drosophila OSNs, where such features are attributed to odorant receptor identity^58^. In mice, further experiments are necessary to understand the determinants of these glutamate dynamics - for example, whether they are determined by the affinity and intensity of an odorant relative to the dynamic range of a particular glomerulus-odorant pair, or instead by properties of odorant sorption in the olfactory epithelium.

Finally, we observed a very close (but not perfect) correspondence between the dynamics of glutamatergic input to MT cells and those of postsynaptic Ca^2+^ signals reflecting MT cell spiking, imaged simultaneously. These results suggest that the dynamics of MT cell spiking patterns closely follow those of glutamatergic input to their glomerulus. Interestingly, we saw no evidence for broadly-organized inhibition surrounding glomeruli receiving excitatory inputs, as predicted by models of center-surround or global inhibition^18,59^. We also saw little evidence for suppression of MT cell output in glomeruli receiving weak excitatory inputs, as predicted from models of feedforward inhibition^11,17,38^. Instead, suppressive MT cell responses were sparse and often co-occurred with a reduction in the glutamate signal to below baseline levels, indicating that MT cell suppression commonly reflects a reduction in excitatory drive. This reduction could be could be due to adaptation of OSN inputs^48^, inhibition at the receptor level^60,61^, or inhibition of ongoing glutamatergic drive from ET cells^11,12^. Overall, the high correspondence between glutamatergic input and MT cell output patterns suggests that transformations of odor representations by OB circuitry- at least in the spatial and temporal domains - may be more limited than previously thought. Instead, inhibitory circuits may function to shape response features such as sensitivity and response gain by acting coincident with excitation. Such a model would be more similar to the function of many inhibitory circuits in cortex, which are reliant on and closely coupled to excitation^62^.

Another surprising aspect of our results is the relatively small impact of blocking postsynaptic activity on glutamate signaling onto MT cells. Studies from OB slices suggest that ET cells provide a large fraction of excitatory drive to MT cells via glutamate spillover from the ET- to - periglomerular cell synapse^16,17,44,50^. Extrapolating to OB circuit function in vivo is difficult, however – for example, glutamate spillover may contribute less to MT cell drive in vivo than in OB slices due to differences in glutamate transporter efficacy or in the dynamics of odorant-evoked glutamate release from OSN inputs. We found that a near-complete blockade of postsynaptic activation, as confirmed with GCaMP reporters expressed in monosynaptically-driven tufted cells, had no impact on the dynamics of inhalation-linked glutamate transients and only modestly impacted the amplitude of transients in response to low odorant concentrations, as well having rather subtle effects on responses across inhalations.

Our results are consistent with some models of ET cell-mediated excitation of MT cells, which propose that the ET cell pathway is most important in regimes of weak OSN input (i.e., low odorant concentrations)^17,43,50^, or that direct OSN inputs are shunted by gap junctions between MT cells^41^, which would not be reflected in our iGluSnFR recordings. Nonetheless, the overall persistence of glutamate signals after postsynaptic blockade, as well as the strong correspondence between glutamate dynamics and MT cell activation, suggests that monosynaptic drive from OSNs is the primary determinant of MT cell response patterns across a large regime of odorant stimulation and natural odor sampling. An alternate possibility is that the disynaptic circuit provides tonic glutamatergic drive to MT cells via ET cell bursting in vivo, allowing for modulation of MT cell excitability by inhibitory circuits. Further use of iGluSnFRs and other transmitter-specific reporters could help disentangle the contributions of excitation and inhibition to shaping dynamic odor representations in vivo.

## MATERIALS AND METHODS

### Animals

Experiments were performed on male and female mice expressing Cre recombinase (Cre) in defined neural populations. Mouse strains used were: Pcdh21-Cre (Tg(Pcdh21-cre)BYoko), Gensat Stock #030952-UCD; OMP-Cre (Tg(Omp-tm4-Cre)Mom), JAX Stock #006668, Tbet-Cre (Tg(Tbx21-cre)1Dlc), JAX Stock #024507, and CCK-IRES-Cre (Tg(CCK-IRES-Cre)Zjh), JAX Stock #012706^22^. Mice ranged from 3-8 months in age. Mice were housed up to 4/cage and kept on a 12/12 h light/dark cycle with food and water available ad libitum. All procedures were carried out following the National Institutes of Health Guide for the Care and Use of Laboratory Animals and were approved by the University of Utah Institutional Animal Care and Use Committee.

### Viral vector expression

Viral vectors were obtained from the University of Pennsylvania Vector Core (AAV1 or 5 serotype, AAV.hSynap-FLEX.iGluSnFR and AAV.hSynap-FLEX.jRGECO1a), Addgene (AAV1 serotype, pAAV.hSynap-FLEX.SF-iGluSnFR.S72A, #106182), HHMI Janelia Campus or Vigene (AAV1 or 5 serotype, pAAV.hSynap-FLEX.SF-iGluSnFR.A184V, pAAV.hSynap-FLEX.SF-iGluSnFR.A184S). Virus injection was done using pressure injections and beveled glass pipettes, as described previously^24,31,63^. For coinjection of jRGECO1a and SF-iGluSnFR.A184S, virus was either diluted 1:10 and injected in separate pipettes through the same craniotomy, or mixed and coinjected via the same pipette. After injection, mice were given carprofen (Rimadyl, S.C., 5 mg/kg; Pfizer) as an analgesic and enrofloxacin (Baytril, I.M., 3 mg/kg; Bayer) as an antibiotic immediately before and 24 hours after surgery. Mice were singly housed after surgery on ventilated racks and used 21-35 days after virus injection. In some mice, viral expression was characterized with post-hoc histology using native fluorescence.

### In vivo two photon imaging

Two-photon imaging in anesthetized mice was performed as described previously^30,63^. Mice were initially anesthetized with pentobarbital (50 - 90 mg/kg) then maintained under isoflurane (0.5 – 1% in O_2_) for data collection. Body temperature and heart rate were maintained at 37 °C and ∼ 400 beats per minute. Mice were double tracheotomized and isoflurane was delivered passively via the tracheotomy tube without contaminating the nasal cavity^64^. Two-photon imaging occurred after removal of the bone overlying the dorsal olfactory bulb and stabilizing the brain surface with agarose and a glass coverslip.

Imaging in awake, head-fixed mice was performed through a chronic imaging window implanted over one dorsal OB. The imaging window consisted of a custom double coverslip with minimum diameter of 1.5 mm. Virus (500-750 nL) was injected at a depth of 250 μm at the time of window implant. Animals were acclimated to the imaging rig for at least one 30-minute session the day prior to imaging. Imaging sessions lasted from 30-60 minutes per day, over the course of several days. The behavior and imaging apparatus has been described previously^30^. Mice were naïve to the odors at their first imaging session. Respiration (sniff) signals were obtained with an external flow sensor (FBAM200DU, First Sensor, AG, Berlin) in front of the animal’s right nostril^65^, or by using a thermistor (MEAS-G22K7MCD419, Measurement Specialties; Hampton, VA) implanted in the right nasal bone^66^.

Imaging was carried out with a two-photon microscope (Sutter Instruments or Neurolabware) coupled to a pulsed Ti:Sapphire laser (Mai Tai HP, Spectra-Physics; or Chameleon Ultra, Coherent) at 920-940 nm and controlled by either Scanimage (Vidrio) or Scanbox (Neurolabware) software. Imaging was performed through a 16X, 0.8 N.A. objective (Nikon) and emitted light detected with GaAsP photomultiplier tubes (Hamamatsu). Fluorescence images were acquired using unidirectional resonance scanning at 15.2 or 15.5 Hz. For SF-iGluSnFR.S72A imaging, bidirectional scanning at 30Hz was utilized to capture faster responses. For dual-color imaging, a second laser (Fidelity-2; Coherent) was utilized to optimally excite jRGECO1a (at 1070nm) and emitted red fluorescence collected with a second PMT, as described previously^24^.

### In vivo pharmacology

In vivo pharmacology was carried out after removing the bone and dura overlying the dorsal olfactory bulb, using protocols described previously^23,46^. Drug solutions (1mM CGP35348 and 0.5 mM NBQX / 1 mM APV) were dissolved in Ringers solution, pre-warmed on a heating block, and applied in bulk to the dorsal OB without the use of agarose or coverslip. For higher concentration experiments, we utilized 2.5 mM NBQX / 5 mM APV dissolved in Ringers. We waited at least 10 minutes after drug application to allow for absorption and temperature equilibration across the tissue. NBQX + APV was applied immediately after CGP35348 without an intervening control wash; due to its extremely slow washout in vivo, we considered CGP35348 to still be present during the subsequent NBQX + APV application. For statistical analysis of pharmacology experiments, all comparisons were done within-animal unless otherwise noted.

### Odorant stimulation

In some experiments, odorants were presented as precise dilutions from saturated vapor (s.v.) in cleaned, desiccated air using a custom olfactometer under computer control, as described previously^30,67^. Odorants were presented for durations ranging from 2 - 8 sec. Clean air was passed across the nostrils in between trials to avoid contribution from extraneous odorants in the environment. Odorants were pre-diluted in solvent (1:10 or 1:25 in mineral oil or medium chain triglyceride oil) to allow for lower final concentrations and then diluted to concentrations ranging from .3% to 1% s.v. Relative increases in concentration were confirmed with a photoionization detector (miniRAE Lite, PGM-7300, RAE Systems) 3 cm away from the flow dilution port. Estimated final concentrations of odorants used ranged from 0.01 – 300 ppm, depending on vapor pressure and s.v. dilution (Table 1). For experiments testing a larger panel of odorants (e.g., Figs. 3 and 5, see Table 1), we used a novel, recently described olfactometer design that allowed for rapid switching between odorants with minimal cross-contamination^68^. Here, odorants were presented for 2 or 3 seconds, in random order from among a bank of 12 odorant cartridges, using 10 second interstimulus intervals. Odorant presentation was as described in a previous publication^68^, using an eductor nozzle for optimal mixing in a carrier stream of filtered air. The end of the eductor was placed 5-7 cm away from the nose. With the configuration used, estimated dilutions of odorant were approximately 1.5% s.v.; odorants were prediluted to achieve relatively sparse activation of dorsal glomeruli^68^. Estimated final concentrations ranged from 0.01 – 16 ppm (Table 1).

### Data Analysis

Image analysis was performed using custom software written in MATLAB (Mathworks). For display, odorant response maps were displayed using ΔF values rather than ΔF/F to minimize noise from nonfluorescent regions. Activity maps were scaled as indicated in the Fig. and were kept to their original resolution (512 x 512 pixels) and smoothed using a Gaussian kernel withσ of 1 pixel. For time-series analysis, regions of interest (ROIs) were chosen manually based on the mean fluorescence image and were further refined based on odorant response ΔF maps, then all pixels averaged within an ROI. All signals were upsampled to 150 Hz for analysis using the MATLAB pchip function. Time-series were typically computed and displayed as ΔF/F, with F defined as the mean fluorescence in the 1-2 sec prior to odorant onset.

For analysis of inhalation-linked dynamics, inhalation-triggered average (ITA) responses were generated by averaging each inhalation (delivered at 0.25 Hz) over a 70 sec odorant presentation (17 inhalations averaged in total). Onset latencies, peak responses, and ITA durations (full-width at half-maximum) were defined as previously^24^. To calculate decay time-constants (Fig. 1), a single exponential was fitted from the peak of the unfiltered ITA trace and extending to the end of the trace. Time-to-peak values were calculated from unfiltered ITAs as the time from the inhalation start to the peak of the response.

For analysis of odorant-evoked dynamics over multiple sniffs, responses to 3 - 8 presentations of odorant were averaged before analysis. Changes in response amplitude over time, or T_2_-T_1_/T_max_ (Fig. 6) were calculated as the difference in amplitude between the peak ΔF following the first (T_1_) and the second-to-last (T_2_) inhalation during odorant presentation, divided by the maximum ΔF during the 4 sec odor presentation.

To analyze response patterns across the 23-odorant panel (i.e., Fig. 3), responses were averaged across 3 - 4 randomized presentations of each odorant. Responses were classified as having significant excitatory and/or suppressive components as follows: First, each averaged response for an ROI-odor pair was low-pass Gaussian filtered at 2 Hz (to test for excitatory responses) or 0.5 Hz (for suppressive responses) and z-scored using a baseline from 0.5 to 2 sec prior to odorant onset, with z defined as the SD of the baseline period concatenated for all 23 odorant responses for each ROI. Peak excitation was measured as 95% of the maximum signal during the 3- or 4-sec odorant presentation; suppressive responses were measured as the 15^th^ percentile of all values in a time-window from odorant onset to 500 ms after odorant offset. We then used a very conservative criterion for significance of z = ±7 SD for identifying significant excitatory or suppressive response components. This cutoff was chosen to yield a false positive rate of approximately 1% (based on visual inspection of a subset of traces).

Glomerular tuning was characterized from response matrices thresholded according to these significance values, using a measure of lifetime sparseness^69^

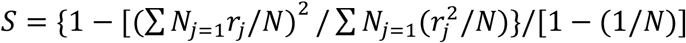

Where *N* is number of odors and *r*_*j*_ is the thresholded z-scored value for a given glomerulus of the response to odor *j*. Sparseness was calculated separately for excitatory and suppressive response components.

Time-dependent decorrelation of odorant response patterns across glomeruli (Figs. 3 and 4) was calculated as previously described ^6^, using all glomeruli in a field of view without thresholding. Each odorant response was binned into five time bins (each of 387 ms) spanning the length of odorant presentation but omitting the first 65 ms to account for delays in odorant onset. Pearson’s correlation coefficient (*r*) was calculated using the response amplitudes across all ROIs in the first time bin as a vector, compared to the same vector taken from each subsequent bin. We calculated an overall correlation time series by computing the mean and s.e.m. for all odorant responses, pooled across all fields of view.

For analysis of inhalation-linked timing of glutamate signals in awake mice, we restricted analysis to inhalations occurring at a minimum period of 200 ms (5 Hz respiration rate). Inhalation timing was detected from the external flow sensor using peak/trough detection and maximal slopes of the flow signal. Inhalation peak was defined as the trough of the external flow signal; inhalation onset was defined as the maximal slope of the transition from exhalation to inhalation. Because inhalation peak timing was a more robust measure, we generated inhalation-triggered average iGluSnFR signals based on inhalation peak. ITAs were generated by averaging the unfiltered optical signal taken from -0.3 sec before to 0.4 sec after inhalation. To avoid confounds of the airflow waveform or measurement method (airflow versus thermistor) in determining absolute ITA response latencies, we report the range of latencies relative to the median of a given set of measurements. Statistical tests were performed either in Origin (OriginLab Corp.), MATLAB (Mathworks) or R (version 3.3.2). Nonparametric tests were used in most cases; parametric tests were used only on datasets determined to be normally distributed. Summary statistics are reported as mean ± standard deviation (SD), unless otherwise stated. All measurement of response parameters was done using analysis code that was independent of treatment or comparison condition.

## ACKNOWLEDGEMENTS

We thank L. Looger and J. Marvin and the GENIE Project from HHMI Janelia Research Campus for sharing SF-iGluSnFR virus. We also thank M. Economo, D. Brunert, Y. Tsuno and F. Fang for initial characterization of iGluSnFRs, G. Vasquez-Opazo, R. Kummer and J. Ball for technical assistance, T. Rust for data analysis software, as well as S. Burton, S. Short, I. Youngstrom, and K. Podgorski for providing helpful feedback and discussion. This work was supported by the NINDS T32NS076067 (to A.K.M) and NIDCD R01DC006441 (to M.W.), R01DC006441-13S1 (to A.K.M), and F32DC015389 (to T.P.E).

## COMPETING INTERESTS

These authors declare no competing financial interests.

## DATA AVAILABILITY

Data underlying the findings of this study are available from the corresponding author upon request.

## CODE AVAILABILITY

Custom code (Matlab) used in data analysis are available from the corresponding author upon request.

## CONTRIBUTIONS

A.K.M and M.W. designed research. A.K.M, T.P.E, and M.W. performed research, A.K.M., T.P.E., and M.W. analyzed data, A.K.M. and M.W. wrote the paper.

## Notes

### Competing Interest Statement

The authors have declared no competing interest.

### Summary of Updates

New data on pharmacological blockade of postsynaptic activity and dual-color imaging of pre- and postsynaptic MT cell signals added. Additional data analyses added, and text and figures revised and reorganized.

