## Supplementary Figures for "Diverse dynamics of glutamatergic input from sensory neurons underlie heterogeneous responses of olfactory bulb outputs *in vivo*"

Supplementary Figure 1

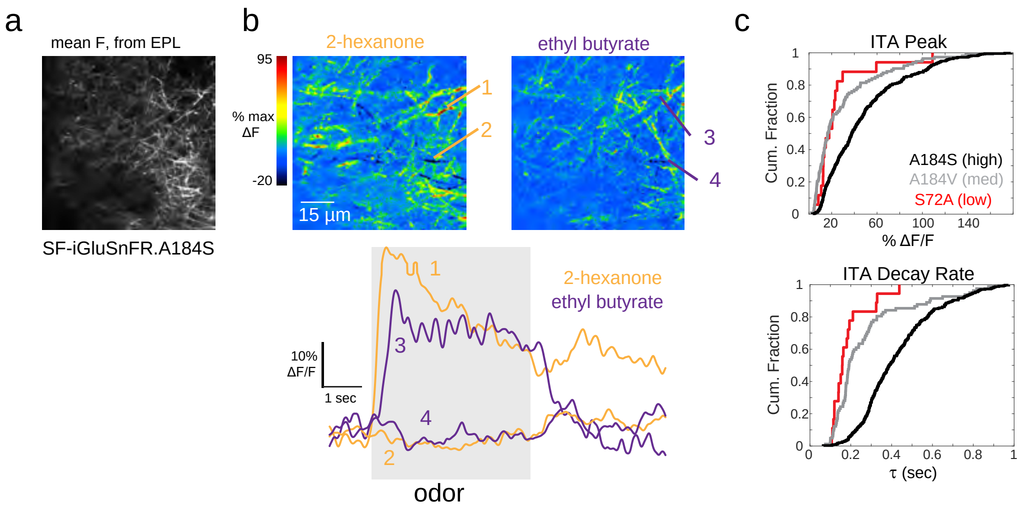


**Supplementary Figure 1. Further characterization of SF-iGluSnFR signals in mitral/tufted cells.**

a. SF-iGluSnFR.A184S expression imaged in vivo from Tbet-positive MT cell lateral dendrites in the superficial external plexiform layer (EPL).
b. Top: Odorant-evoked response maps imaged from the field of view in (a), showing responses to two odorants, 2-hexanone and ethyl butyrate. Bottom: Traces showing stronger excitatory and weak suppressive SF-iGluSnFR signals on different dendrites, evoked by each odorant.
c. Cumulative fraction of peak ITA response magnitudes (top) and decay rates (bottom) across the population of significantly-responding glomerulus-odor pairs measured for each SF-variant. Note: A184S is the high affinity variant, A184V is the medium affinity variant, and S72A is the low affinity variant. Maximal response amplitudes (90th percentile of max ΔF/F across all glomerulus-odor pairs) were 59% for S72A (n = 18 glomerulus-odor pairs, 2 mice); 74% for A184V (n = 117, 2 mice); and 104% for A184S (n = 438, 5 mice). Minimal decay rates (25th percentile across all glomerulus-odor pairs) were 120 ms for S72A (n = 18 glomerulus-odor pairs from 2 mice), 162 ms for A184V (n=91, 2 mice) and 297 ms for A184S (n = 370, 5 mice).

Supplementary Figure 2

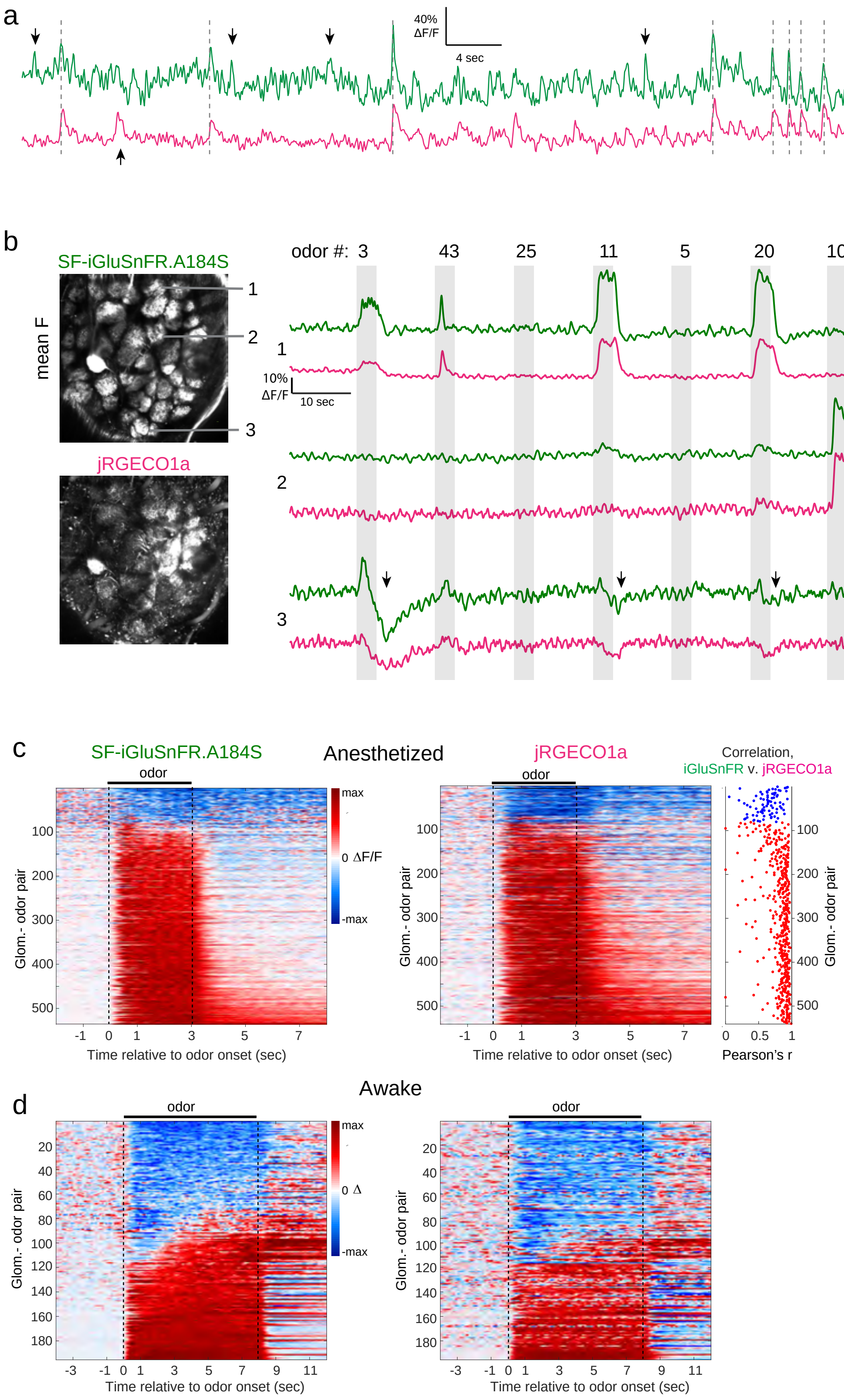

**Supplementary Figure 2. High correspondence between glomerular glutamate and calcium signals imaged simultaneously with dual-color two-photon imaging.**

a. Traces showing SF-iGluSnFR and jRGECO1a signals imaged simultaneously from a glomerulus, with high correspondence between spontaneously-occurring transients. Vertical lines mark transients seen in both signals; downward arrows mark transients seen in the green (A184S) but not red (jRGECO1a) channels; upward arrows mark transients seen in the red but not the green channel.
b. Left: Mean fluorescence images for SF-iGluSnFR.A184S and jRGECO1a from the same field of view. Right: Traces showing continuous dual-color recording across 10 consecutive odorant presentations, taken from three glomeruli (locations shown in mean fluorescence images at left), showing high correspondence in SF-iGluSnFR.A184S and jRGECO1a signals despite diverse response patterns to different odorants. ROI 3 (lower traces) shows clear suppressive components in both signals. Downward arrows denote a biphasic response in the SF-iGLuSnFR signal that is not present in the jRGECO1a signal.
c. 'Waterfall' plots showing Top: Anesthetized and Bottom: Awake waterfall plots showing time-course of SF-iGluSnFR.A184S and jRGECO1a signals for all significantly-responding glomerulus-odor pairs imaged from anesthetized mice (5 fields of view in 3 mice). Each row is z-scored and scaled to its maximal odorant-evoked deviation from baseline; row order is sorted by the summed SF-iGluSnFR.A184S response across the odorant presentation window (sort order is the same for SF-iGluSnFR and jRGECO1a signals). Right: Correlation coefficients (Pearson's r) of the time-course of the odorant response in each channel (SF-iGluSnFR and jRGECO1a), shown for every glomerulus-odor pair. Correlations are generally high, but are most variable for suppressive responses (blue).
d. 'Waterfall' plots, displayed as in (c) but for all significantly responding glomerulus-odorant pairs in awake mice (2 fields of view, two mice).

Supplementary Figure 3

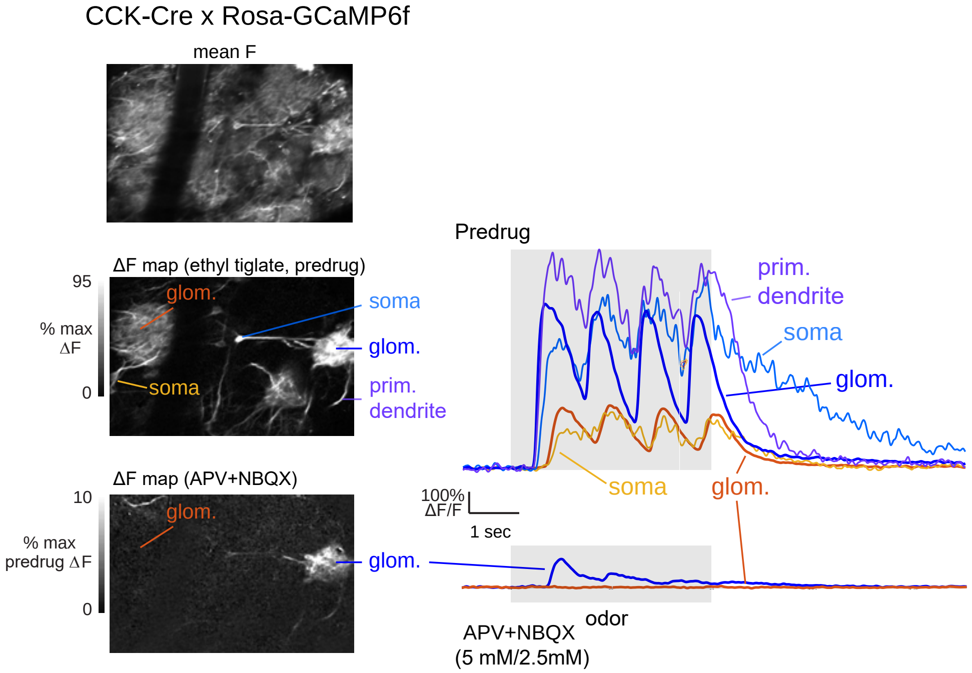


**Supplementary Figure 3. Pharmacological blockade of calcium signals in dendritic and somatic compartments of tufted cells.**

Left: Images show mean fluorescence (top) and ΔF odorant response maps for GCaMP6f signals imaged in a CCK-IRES-Cre : Rosa-GCaMP6f cross. Shown are response maps to the same odorant (ethyl tiglate) before and after APV+NBQX (5 mM/2.5 mM) application. Post-drug response map is scaled to 10% of the predrug response to emphasize small-amplitude persistent response in one of the three activated glomeruli.

Right: Top set of traces show time-course of GCaMP6f signal from the neuropile of two glomeruli (blue, orange traces) and, for each, the soma of a tufted cell innervating each glomerulus. For the glomerulus on the right, the signal from the primary dendrite of a second tufted cell innervating the same glomerulus is also shown. Near-synchronous, inhalation-driven transients are seen in all compartments, with a slightly slower rise and slower decay in the somata. Latency differences between the two glomeruli are also present in their respective cells' somata. Lower traces show the GCaMP6f signal from the neuropile of the two glomeruli, after APV+NBQX application. Note the small but delayed signal remaining in the right glomerulus, and complete blockade in the left glomerulus. Traces from soma and primary dendrite are not shown due to the very poor signal-to-noise ratio of the remaining signal.

Supplementary Figure 4

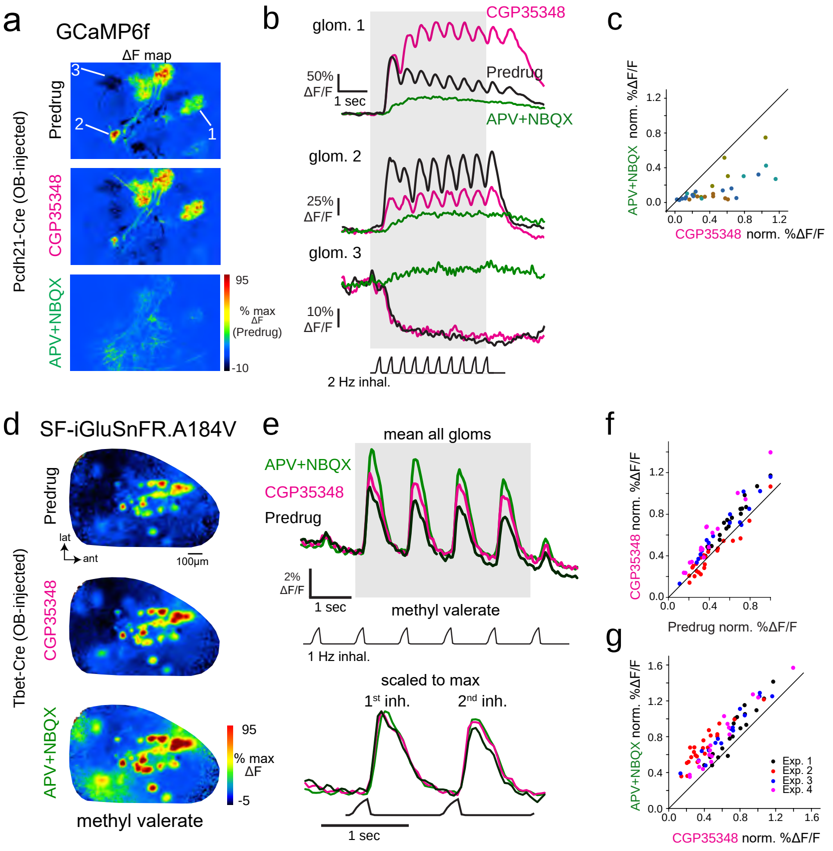


**Supplementary Figure 4. Further characterization of ionotropic glutamate receptor blockade on MT cell postsynaptic responses and glutamate transients.**

a. Odorant-evoked Ca2+ signals imaged with GCaMP6f expressed in Pcdh21-+ MT cells. Maps show responses during predrug, after application of CGP35348, and after subsequent application of APV+N-BQX. Pseudocolor scale normalized to predrug response levels. Note sharp reduction of excitatory responses and elimination of suppressive responses.

b. Response traces for the three glomeruli indicated in (a), (2 are excited, 1 is suppressed) across each condition.

c. Scatter plot comparing GCaMP6f responses across CGP35348 and subsequent APV+NBQX conditions (4 mice). Points show peak response amplitudes to the first inhalation of odorant plotted and normalized as in (c). Each color indicates glomeruli from a different mouse. APV+NBQX/CGP35348 ratio: 0.37 ± 0.06, mean ± s.e.m. of medians in 4 mice, 30 glomeruli; Repeated Measures ANOVA, F1,45 = 10.8, p = 0.002.).

d. Odorant-evoked iGluSnFR response maps imaged across the dorsal OB with epifluorescence in PCdh21+ MT cells, showing glomerular responses before and after application of CGP35348 and APV+NBQX. Maps are scaled to the maximum of predrug levels. Note persistence of glomerular pattern after drug application.

e. Traces showing iGluSnFR signal averaged across responsive glomeruli (same preparation as in (a)) before and after application of CGP35348, followed by APV+NBQX. Lower traces show mean responses over the first two inhalations scaled to the same peak value, indicating little to no change in time-course of the response.

f. Comparison of peak iGluSnFR responses to the first odorant inhalation across predrug and CGP35348 conditions. Each point indicates a glomerulus; each color is from a separate mouse. Points are normalized to the maximum responding glomerulus in the predrug condition for each mouse. Summary statistics: Wilcoxon signed ranks paired test with Bonferroni correction: mouse 1, n = 19, Z = -3.80, p = 5.6 x 10-4; mouse 2, n = 18, Z = 1.22, p = 0.8; mouse 3, n = 17, Z = -3.50, p = 0.002; mouse 4, n = 14, Z = -3.26, p = 0.004).

g. Comparison of iGluSnFR responses across CGP35348 and APV+NBQX conditions; same mice and normalization as in (c). Responses increased significantly in all 4 mice (Wilcoxon signed-ranks; inh. 1: p=0.002, p = 2.6 x 10-4, p = 0.001, p = 0.004).
